# ROR and RYK extracellular region structures suggest that receptor tyrosine kinases have distinct WNT-recognition modes

**DOI:** 10.1101/2021.04.29.442059

**Authors:** Fumin Shi, Jeannine M. Mendrola, Joshua B. Sheetz, Neo Wu, Anselm Sommer, Kelsey F. Speer, Jasprina N. Noordermeer, Zhong-Yuan Kan, Kay Perry, S. Walter Englander, Steven E. Stayrook, Lee G. Fradkin, Mark A. Lemmon

## Abstract

WNTs play key roles in development and disease, by binding both Frizzled (FZD) seven-pass transmembrane receptors and numerous co-receptors that include the ROR and RYK receptor tyrosine kinases (RTKs). We describe crystal structures and WNT-binding characteristics of extracellular regions from the *Drosophila* ROR and RYK orthologs Nrk (neurospecific receptor tyrosine kinase) and Derailed-2 (Drl-2). RORs bind WNTs though a FZD-related cysteine-rich domain (CRD), and RYKs through a WNT-inhibitory factor (WIF) domain. Our structures suggest that neither the Nrk CRD nor the Drl-2 WIF domain can accommodate the acyl chain typically attached to WNTs. The Nrk CRD contains a deeply buried bound fatty acid, unlikely to be exchangeable with a WNT acyl chain. The Drl-2 WIF domain lacks the lipid-binding site seen in WIF-1. We also show that DWnt-5, which regulates *Drosophila* ROR and RYK orthologs, lacks an acyl chain. Together with analysis of WNT/receptor interaction sites, these structures provide new insight into how WNTs recruit their RTK co-receptors into signaling complexes.

## INTRODUCTION

WNTs play diverse roles in development, adult stem cell renewal and tissue homeostasis (Nusse and Clevers, 2017), and are represented in humans by 19 different genes that have different functions (Miller, 2012). Given their roles in a variety of developmental processes and numerous diseases – from cancer to developmental defects to degenerative diseases (Nusse and Clevers, 2017) – there is a great deal of interest in understanding mechanisms of WNT signaling in order to develop approaches to modulate it therapeutically. The best known receptors for WNT are called Frizzleds (FZDs), members of the F class of G protein-coupled receptors (GPCRs), which have an extracellular cysteine-rich domain (CRD) of ∼120 aa that directly binds WNTs (MacDonald and He, 2012). The details of WNT binding to the extracellular CRD of FZD8 have been visualized in crystal structures (Hirai et al., 2019; Janda et al., 2012), with interactions between a WNT-associated acyl chain and a hydrophobic channel on the CRD surface being key. This acyl chain recognition appears to promote FZD dimerization (Hirai et al., 2019) through a characteristic interface also utilized by FZD CRDs bound to isolated fatty acids (DeBruine et al., 2017; Nile and Hannoush, 2019; Nile et al., 2017). Beyond possibly promoting FZD homodimerization, bound WNTs appear to function primarily as cross-linkers to bridge FZDs to co-receptors such as LRP5/6 (Janda et al., 2017; Tao et al., 2019) – without directly altering the conformation of the transmembrane region of the FZD protein (Tsutsumi et al., 2020).

Four of 20 families of human receptor tyrosine kinases (RTKs), which control many different cellular processes (Lemmon and Schlessinger, 2010), are now known also to be receptors or co-receptors for WNTs (Green et al., 2014; Niehrs, 2012; Roy et al., 2018). These are the ROR family (for receptor tyrosine kinase-like <u>o</u>rphan receptor), RYK (for receptor t<u>y</u>rosine <u>k</u>inase-related tyrosine kinase), PTK7 (for <u>p</u>rotein tyrosine <u>k</u>inase-<u>7</u>), and MuSK (for <u>mu</u>scle-<u>s</u>pecific receptor tyrosine <u>k</u>inase), which have all been shown to be involved in multiple aspects of WNT signaling (Fradkin et al., 2010; Green et al., 2008; Jing et al., 2009; Lhoumeau et al., 2011). Consistent with a role as direct WNT receptors, the RORs and MuSK contain an extracellular cysteine-rich domain (CRD) related to that seen in FZDs. RYK instead contains a WIF (for <u>W</u>NT-Inhibitory <u>F</u>actor) domain in its extracellular region, again implicating it in WNT binding (Hsieh et al., 1999). Interestingly, the RORs, RYK, and PTK7 also stand out among RTKs by having pseudokinases in their intracellular regions (Mendrola et al., 2013; Sheetz et al., 2020). Thus, this group of WNT receptors is likely to differ from both the FZD family and canonical RTKs in their signaling mechanisms.

Given the importance of ligand-induced dimerization in regulation of canonical RTKs (Lemmon and Schlessinger, 2010), we were interested in understanding how WNTs engage ROR and RYK family members. We were also interested in assessing the importance of WNT acylation for ROR and RYK regulation, since sequence alignments have suggested that ROR family CRDs do not maintain the hydrophobic channel seen in FZD CRDs (Janda and Garcia, 2015). Moreover, we have found in previous studies that WNT acylation is not always required for signaling (Speer et al., 2019). Combining structural studies of extracellular regions (ECRs) from ROR and RYK family members with ligand-binding studies and other analyses, we show here that the requirements for WNT-attached fatty acids are quite different for binding of WNTs to these RTK co-receptors than for binding to canonical FZDs. We also find that WNT binding does not appear to induce RTK homodimerization. Our results suggest a model in which WNTs cross-link RTK pseudokinase co-receptors into signaling complexes with FZDs, within which they might allosterically regulate other key components.

## RESULTS AND DISCUSSION

In efforts to investigate how WNT-regulated RTKs are engaged by their ligands, we found that extracellular regions (ECRs) from *Drosophila melanogaster* ROR and RYK family members behaved best in biophysical and crystallization studies. The *Drosophila* ROR orthologs dRor and Nrk (for neurospecific receptor tyrosine kinase; also called dRor2) are expressed specifically in the fly nervous system during embryogenesis (Oishi et al., 1997; Ripp et al., 2018). The three *Drosophila* RYK family members are Derailed (Drl), Drl-2 and Doughnut on 2 (Dnt). They play key roles in neuronal pathway selection (Callahan et al., 1995; Inaki et al., 2007) and muscle attachment site targeting (Callahan et al., 1996; Lahaye et al., 2012; Yoshikawa et al., 2001), as well as other aspects of fly nervous system function, including synaptic growth (Liebl et al., 2008), olfactory system patterning (Hing et al., 2020; Sakurai et al., 2009; Wu et al., 2014), mushroom body development (Reynaud et al., 2015), and peripheral nervous system wiring (Yasunaga et al., 2015). Both ROR and RYK families of receptors are thought to be regulated by the *Drosophila* WNT-5 ortholog, DWnt-5 (Lahaye et al., 2012; Ripp et al., 2018; Wouda et al., 2008; Yoshikawa et al., 2003).

### Structure of the Nrk ECR

We first determined the crystal structure of the Nrk ECR (sNrk) to 1.75 Å resolution (Table 1). In *Drosophila*, the dRor and Nrk ECRs contain a FZD-like cysteine-rich domain (CRD) followed by a kringle (Kr) domain (Figure 1A), but lack the amino-terminal immunoglobulin (Ig)-like domain seen in human ROR1 and ROR2 (Roy et al., 2018). The linker between the CRD and Kr domains is well ordered in sNrk, and the two domains together appear to form a single unit (Figure 1B and C) – with 1160 Å^2^ of surface area buried between the two domains (580 Å^2^ each). Parallel small angle X-ray scattering (SAXS) studies of the human ROR2 ECR (Figure S1A-D) indicate that the amino-terminal Ig-like domain in humans extends this globular structure into a longer rigid rod. As shown in Figure S1E, the Kr domain of sNrk overlays very well (RMSD < 2 Å, for all atoms) with those from human ROR1 (Qi et al., 2018) and ROR2 (Goydel et al., 2020), determined previously as fragments bound to potential therapeutic antibodies. The Nrk CRD closely resembles the CRDs seen in FZDs (Dann et al., 2001; Nile and Hannoush, 2019), Smoothened (Nachtergaele et al., 2013), and MuSK (Stiegler et al., 2009). The domain is mostly α-helical, and is stabilized by 5 disulfide bonds that are very similar in location (and identical in connectivity) in all but the Smoothened CRD (Nachtergaele et al., 2013) – which lacks one disulfide bond. An amino-terminal hairpin-like structure precedes the first α-helix (α1/2), and a β-hairpin (containing strands β3 and β4 in Nrk) connects α3 and α4 (using the secondary structure element numbering of Dann et al., 2001). In a Dali search (Holm, 2020) the Nrk CRD is most similar to that from rat MuSK (Figures 2A,B), with which it shares 30% sequence identity (Xu and Nusse, 1998). The Nrk and MuSK CRDs differ in significant ways from those in FZDs, among which the closest homolog of Nrk is FZD4, with 17% sequence identity (Figure S2).

**FIGURE 1.**
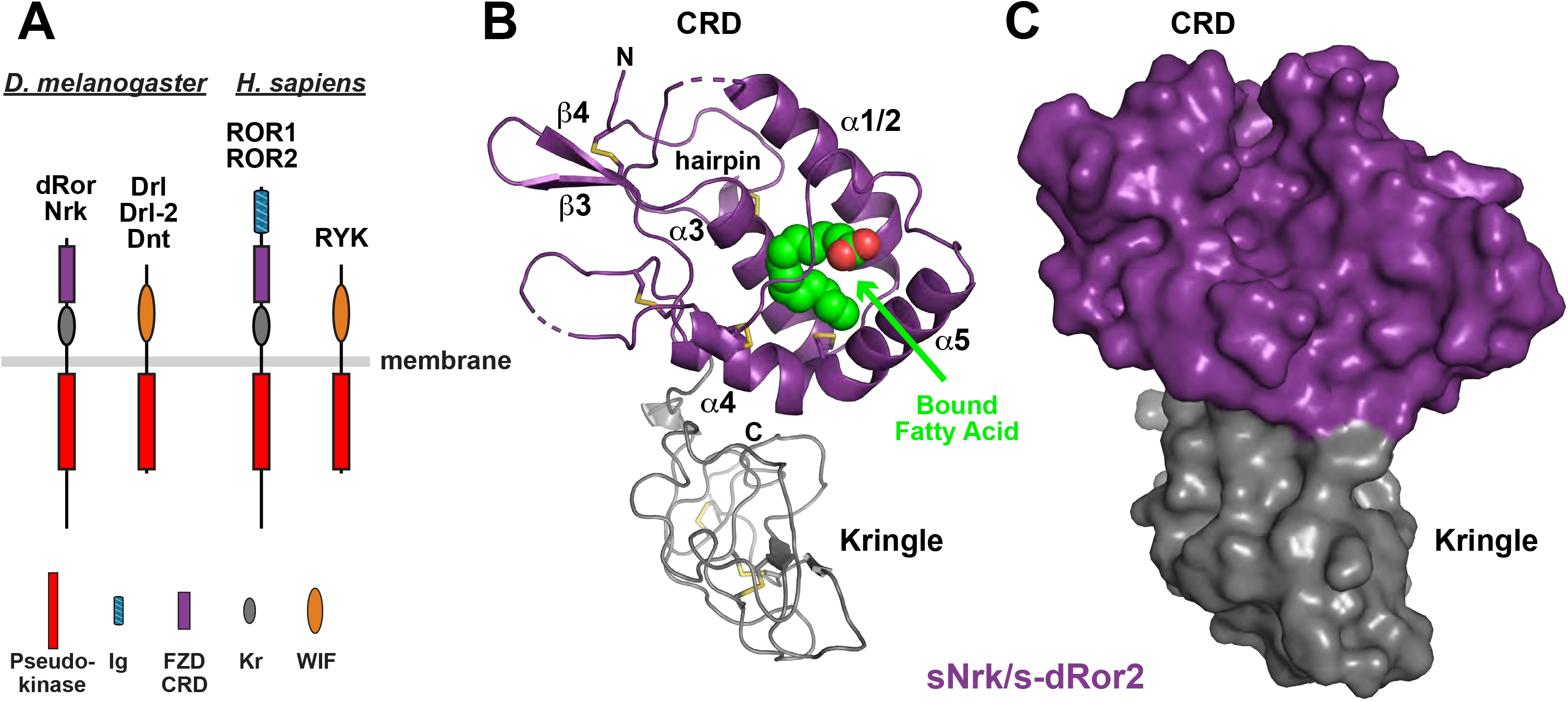
Structure of a ROR family ECR. **(A)** Domain composition of ROR and RYK/Drl family RTKs from *D. melanogaster* (left) and *H. sapiens* (right). The membrane is depicted as a horizontal gray line. As shown in legend, the pseudokinase domain is colored red, immunoglobulin-like domain is blue, FZD-related CRD is purple, Kringle domain is grey, and WIF domain orange. Note that there are three RYK orthologs in *D. melanogaster*, but only one in humans. **(B)** Cartoon representation of the sNrk/s-dRor2 structure, with the CRD colored purple and Kringle domain colored grey. Secondary structure elements are labeled in the CRD only – using the designation introduced by the Leahy lab (Dann et al., 2001) – and the bound fatty acid molecule is shown as green and red spheres. Disulfides are shown as sticks, and the N-terminal hairpin is marked. **(C)** Surface representation of sNrk/s-dRor2, colored as in B. Note that burial of the bound fatty acid molecule causes it not to be visible in this representation. See also Figure S1.

**FIGURE 2.**
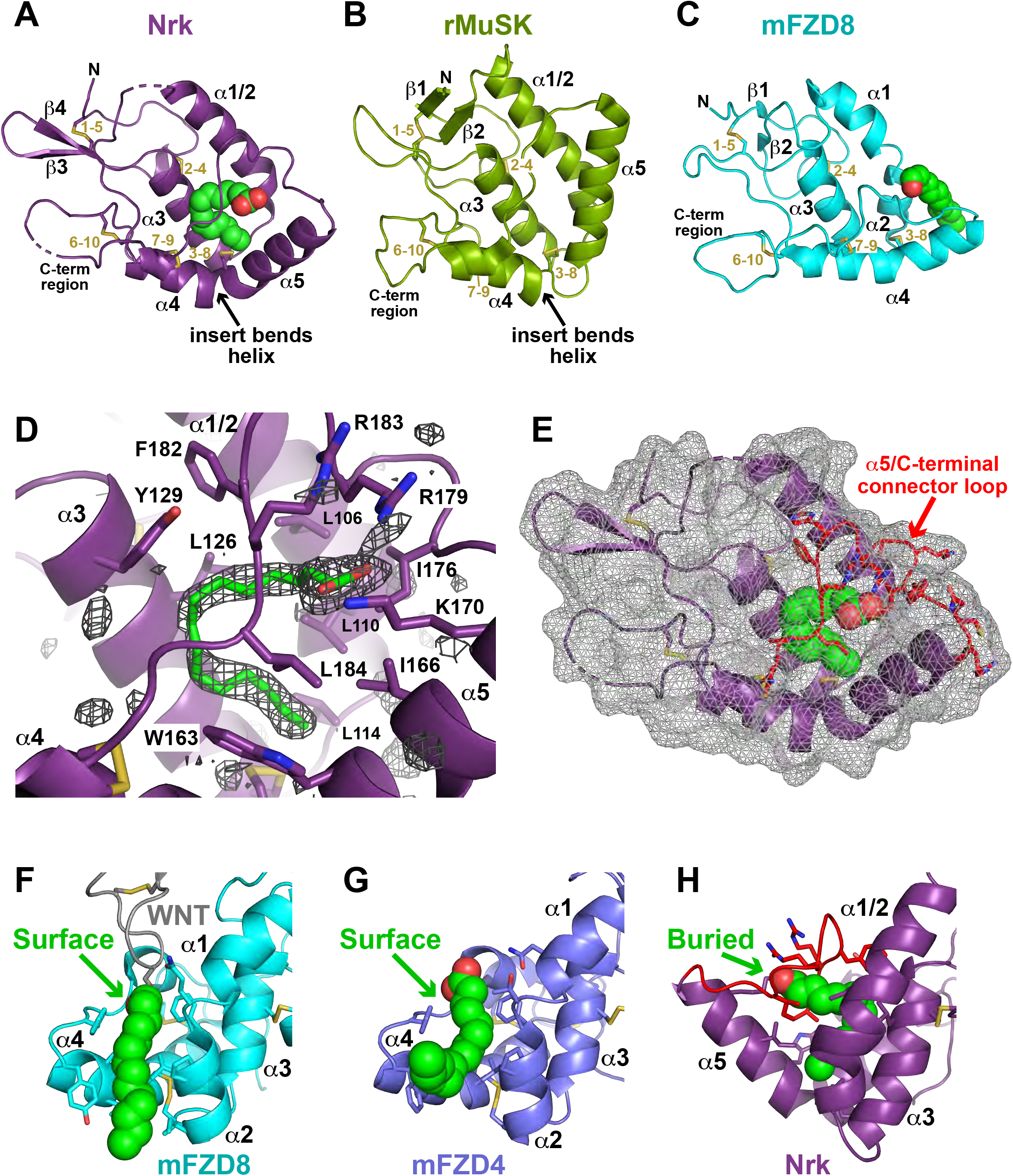
Comparison of ROR and FZD family CRDs and modes of fatty acid binding. **(A)** Cartoon representation of the sNrk CRD in the same orientation as in Figure 1B, with secondary structure elements marked. Disulfide bonds are numbered (in gold) for the cysteine order, and the bound fatty acid is shown in spheres. The bend in helix α4, yielding the C-terminal α5 helix, is marked – as is the C-terminal region. **(B)** Cartoon representation of CRD from rat MuSK (PDBID: 3HKL), in the same orientation as used for Nrk in A, and labeled similarly. Note the absence of bound lipid, but retention of the bend in helix α4. **(C)** Cartoon representation of CRD from mouse FZD8 (PDBID: 1IJY), in the same orientation as used for Nrk in A, and labeled similarly. A bound palmitoleic acid is shown in green spheres based on its position in the xWnt8/mFZD8 complex (PDBID: 4F0A). **(D)** Unbiased |F_o_|-|F_c_| Polder omit map, contoured at 3 σ, of the region surrounding the site at which the palmitic acid moiety is seen bound to the Nrk CRD. The modeled fatty acid is shown in green sticks, and adjacent secondary structure elements and contacting side-chains are labeled. Note that the basic side-chains from K170, R179, and R183 ‘clamp’ the carboxylate of the fatty acid in position. **(E)** The Nrk CRD is shown with the same orientation as used in A, but with the surface shown as transparent mesh. This representation reveals that the bound fatty acid is completely inaccessible from the domain’s surface – and thus completely buried. The α5/C-terminal connector, which clamps the fatty acid in position, is colored red. **(F)** Illustration of how the mFZD8 CRD (colored cyan) engages the fatty acid attached to xWnt8. The acyl chain of fatty acid covalently attached to xWnt8 is depicted by green spheres (xWnt8 is grey), and lies in a surface channel formed by helices α2 and α4 (Janda et al., 2012) as described in the text. **(G)** Binding of a free fatty acid (green spheres) to the mFZD4 CRD (slate blue), shown in the same orientation as in F, from PDBID: 5UWG. As described (DeBruine et al., 2017), the same surface channel formed by helices α2 and α4 accommodates the acyl chain in this case. **(H)** Fatty acid binding to the Nrk CRD, shown in the same orientation as in F and G, illustrating that the position of the fatty acid binding site is different. The bound fatty acid is fully buried, and CRD structural changes occlude the channel between α2 and α4 as described in the text. See also Figure S2.

**TABLE 1.**
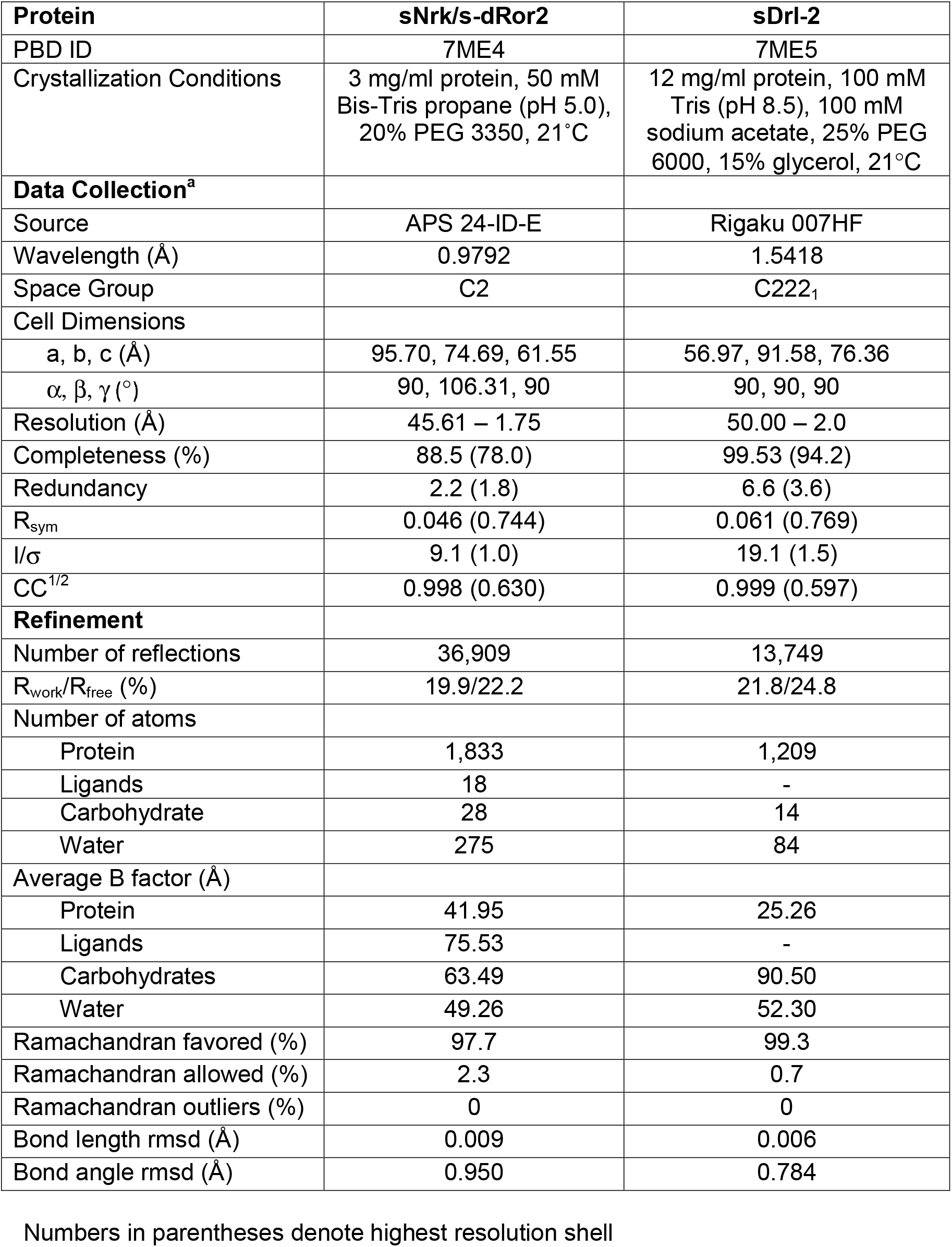
Crystallization Conditions, Data Collection, and Refinement Statistics.

### Key differences between the Nrk and FZD family CRDs

The first notable difference between ROR/MuSK and FZD CRDs is in the most N-terminal helical segment, which begins with a single helix (named α1/2 here; Fig 2A,B) in Nrk and MuSK (Stiegler et al., 2009), but is split into two helices (α1 and α2) in FZD8 (Figure 2C) and other FZD CRDs (Dann et al., 2001). The second difference involves a single-residue insertion before the eighth cysteine of ROR family and MuSK CRDs (Q157 in Nrk – see Figure S2A), which alters α4 in a significant way. In FZD8, the seventh and eight cysteines are both in helix α4, and form disulfide bonds with cysteines 9 and 3 respectively (Figure 2C), after which α4 continues in the same direction. Inserting a residue before cysteine 8 causes a bulge and a bend in this helix for Nrk and MuSK (marked in Figures 2A,B) – effectively breaking the helix in two (helices α4 and α5), with important consequences for acyl chain binding as described below. Helix α5 is also extended in Nrk, MuSK, and other RORs (Figure S2A), and the loop that connects α5 to the C-terminal region of the domain is significantly longer in ROR and MuSK CRDs than the loop that connects α4 to this region in FZD CRDs (Figure S2A).

### The Nrk CRD binds a fatty acid molecule

The Nrk CRD has a bound fatty acid molecule (green in Figure 1B) that was not added during purification or crystallization. It is located within a large (∼855 Å^3^) buried internal cavity between the CRD α-helices. The electron density is consistent with a 16-carbon fatty acid (Figure 2D), but we were not able to unambiguously determine the degree of unsaturation from our structure or by mass spectrometry. Considering palmitic (C16:0) and palmitoleic (C16:1) acids as the likely ligands, the carbon-carbon bond angles at the Δ position in our refined model were better fit with a freely rotating saturated palmitoyl chain, although a palmitoleoyl chain cannot be excluded. Interestingly, no similar internal cavity can be identified between the α-helices of other published CRD structures – including that from MuSK.

### The Nrk CRD fatty acid is deeply buried in the domain

Although fatty acids have been seen bound to several CRDs (Nile and Hannoush, 2019), the mode of binding to the Nrk CRD is quite different than seen for FZD and Smoothened CRDs (Byrne et al., 2016; Nile and Hannoush, 2019). The fatty acid molecule is completely buried in the middle of the Nrk CRD – contacting side-chains from the middle of all four helices (α1/2, α3, α4, and α5; see Figure S2A), which together fully enshroud its ‘U’/’C’-shaped aliphatic region. The carboxylate group of the fatty acid is also buried beneath the loop/insert that follows α5 in Nrk and connects it to the C-terminal part of the CRD (red in Figure 2E) – with the basic side-chains of K170, R179, and R183 ‘clamping’ the carboxylate in place (Figure 2D). As a result, the bound fatty acid appears completely inaccessible from the surface of the Nrk CRD (Figures 2E and S2B) – and fully buried.

This complete burial of the bound fatty acid in the Nrk CRD contrasts with the peripheral accommodation of fatty acids by FZD-family CRDs. Structures of Xenopus Wnt8 (Janda et al., 2012) or human WNT3 (Hirai et al., 2019) bound to the mFZD8 CRD showed that the WNT-attached palmitoleic acid lies in a hydrophobic channel on the CRD surface (Figures 2F, S2C). Interestingly, the same surface-accessible channel was also occupied by a fatty acid molecule in structures of isolated CRDs from FZD4 (Figure 2G), FZD5 and FZD7 (DeBruine et al., 2017; Nile et al., 2017) – even when no lipid was added during purification or crystallization. Moreover, the same surface-lying channel is utilized by human FZD2 to bind a fatty acid (non-covalently) associated with the *Clostridium difficile* toxin B (TcdB) protein (Chen et al., 2018), and by the related Smoothened CRD in binding to cholesterol (Byrne et al., 2016). This channel is formed largely by helices α2 and α4 in mFZD8 and mFZD4 (Figures 2F,G), and the side-chains involved are well conserved across FZD family CRDs (Figure S2A). As shown in Figure 2H, the merging of helices α1 and α2 in Nrk, without a bend between them, causes the end of helix α1/2 to occlude the hydrophobic channel that is seen in FZD CRDs. In addition, the shortening of helix α4 and its projection in a new direction as helix α5 removes the left-hand wall of the channel. Interestingly, the residues that contact the bound fatty acid largely appear similar in position in sequence alignments of ROR family and FZD CRDs (Figure S2A). However, whereas they lie in the channel between α2 and α4 in FZDs, they are instead in the C-terminal part of α1/2 and in α5 respectively in Nrk – relocated in the structure because of the changes summarized above. The bend between α4 and α5, and the new direction of α5 effectively create a new, deeper, fatty acid binding site in Nrk that is also reached by side-chains from α3 (Figures 2D and S2B). In other words, the conserved fatty acid binding site is effectively relocated from the surface to the core of the domain in Nrk (Figures S2B,C), and presumably also in human RORs based on their sequence similarity.

As shown in Figure S2, seven of the 19 side-chains that directly contact the bound fatty acid in Nrk are identical in human ROR1 and/or ROR2 (L114, A122, L126, Y129, L146, L184, P185), and seven are similar (L/F, D/E, W/Y, I/A, and F/L substitutions). Only four are replaced with different residue types (L106, L110, M125, and T154 – replaced by threonine, serine and leucine respectively). These similarities argue that the CRDs of human ROR1 and ROR2 ECRs are likely to bind fatty acids in a related way, although we were unable to identify lipids bound to human ROR ECRs secreted from Sf9 cells using mass spectrometry. The depth of the fatty acid bound in the CRD, and its inaccessibility from the domain’s surface suggest that the lipid plays a co-factor role in this case, rather than playing a part in WNT protein binding.

Interestingly, the side-chains involved in fatty acid binding to Nrk and FZD CRDs tend to be slightly less well conserved in MuSK, and the CASTp server (Tian et al., 2018) shows no significant cavities in the published MuSK CRD structure (Stiegler et al., 2009) that could accommodate an acyl chain of the type bound to sNrk or FZD CRDs.

### The Nrk CRD does not form dimers seen for FZD CRDs

One consequence of the fact that the bound fatty acid is fully buried in the Nrk CRD (and likely CRDs from ROR1/2) is that it cannot mediate CRD dimerization as reported for FZD CRDs (Nile and Hannoush, 2019). Indeed, both sNrk and the hROR2 ECR were monomeric by size exclusion chromatography and in sedimentation equilibrium analytical ultracentrifugation studies. Moreover, the only dimer interface that buries more than 400 Å^2^ in the sNrk crystals is mediated primarily by the Kr domain (Figure S1F). By contrast, the isolated CRDs from FZDs 5, 7, and 8 all crystallized with a similar dimeric relationship that involves a characteristic α-helical dimer (Nile and Hannoush, 2019) also seen for the WNT3-bound mFZD8 CRD (Hirai et al., 2019). These symmetric CRD dimers are mediated in part by α1/α1 and α4/α4 interactions (Figure S2D) – both helices that are altered structurally in ROR family CRDs. In addition, the hydrophobic channel formed between α2 and α4 in one FZD CRD molecule (Figure S2C) is apposed to the equivalent channel in its neighbor, so that a fatty acid bound in this channel can stabilize the dimer – spanning the interface as shown in Figure S2D (Hirai et al., 2019; Nile and Hannoush, 2019; Nile et al., 2017). Whereas fatty acid binding to peripheral locations on FZD family (and possibly Smoothened) CRDs appear to promote homo- or hetero-typic protein-protein interactions (DeBruine et al., 2017; Hirai et al., 2019; Nile and Hannoush, 2019), the structural features of ROR family CRDs do not appear to support this function.

### A RYK/Drl WIF domain has no acyl chain binding site

We also determined the sDrl-2 structure to 2 Å resolution (Table 1). As illustrated in Figure 1A, RYK/Drl family proteins contain a WNT-Inhibitory Factor (WIF) domain in their extracellular region (Callahan et al., 1995; Roy et al., 2018), which takes up almost the entire ECR (residues 26-161 of ∼180 in Drl-2). As shown in Figure 3A, the Drl-2 WIF domain forms a 9-stranded β-sandwich with two short α-helices (α1 and α2) that are ‘presented’ at one of the splayed corners of the β-sandwich (Chothia, 1984). The Drl-2 WIF domain is very similar to the corresponding domain in WIF-1 (Liepinsh et al., 2006; Malinauskas et al., 2011) that is shown in Figure 3B.

**FIGURE 3.**
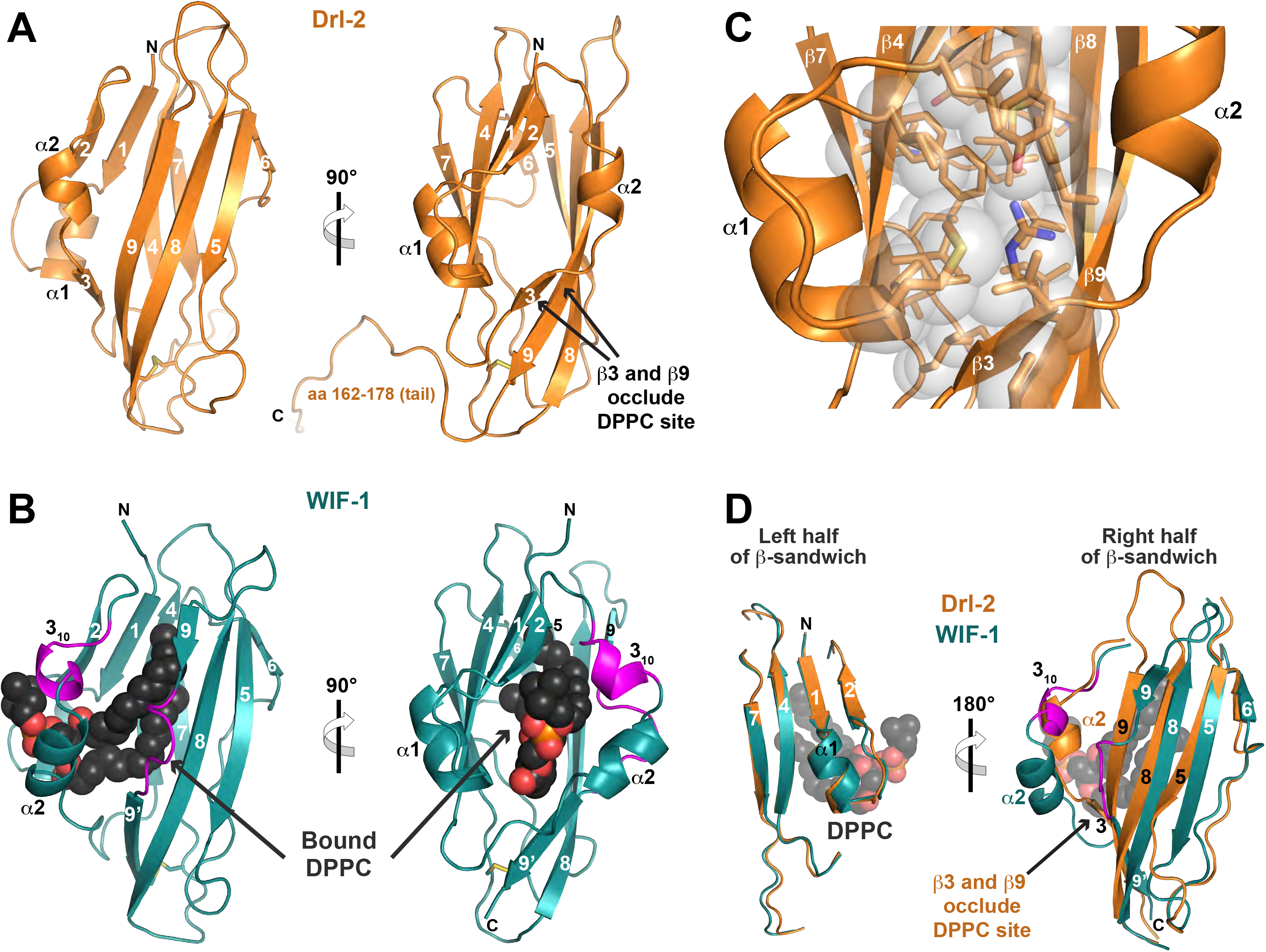
RYK family WIF domain structure and comparison with WIF-1. **(A)** The structure of the complete Drl-2 ECR (colored orange) is shown in cartoon representation in two orthogonal views. Secondary structure elements are marked – using the designation introduced by Liepinsh et al. (2006). The WIF domain ends at around residue 161 (see Figure S3), and the remainder of the ECR (aa 162-182) is involved in crystal packing. **(B)** Cartoon view of the WIF domain from human WIF-1 (PDBID: 2YGN), overlaid and shown in the same orientations as in A (Malinauskas et al., 2011). The WIF domain is colored deep teal, and the bound DPPC molecule is shown as black spheres (red and orange for phosphates). The two regions described as inserts in the WIF-1 WIF domain (a 3_10_ helix and insert in β9) compared with that in Drl-2 are colored magenta. **(C)** Closer view of the hydrophobic core of the Drl-2 WIF domain, using the same orientation as the right-hand side of A, showing that it is well packed, with no cavity capable of accommodating a lipid molecule. **(D)** Overlay of the Drl-2 and WIF-1 WIF domains in two halves as described in the text. The half of the sandwich including β1, β2, β4, β7, and α1 overlay very well. The other half shows more deviations, in particular around α2 (where the magenta insert forms a 3_10_ helix) and in β9 (where a magenta loop/bulge is seen). These changes allow DPPC to bind the WIF-1 WIF domain, whereas the longer (and straight) β9 – along with β3 – occludes the potential lipid-binding site in Drl-2 (and likely other RYK family members). See also Figure S3.

An important key difference between the WIF domains from Drl-2 and WIF-1 is in their ability to accommodate a bound lipid in the middle of the β-sandwich. NMR studies of the WIF domain from WIF-1 (Liepinsh et al., 2006) suggested the existence of a significant internal cavity, suggested by docking studies to accommodate a fatty acid (Malinauskas, 2008). The subsequently determined WIF-1 crystal structure (Malinauskas et al., 2011) revealed that its WIF domain binds a dipalmitoylphosphatidylcholine (DPPC) molecule in the middle of the β-sandwich (Figure 3B). The aliphatic portion of the bound DPPC is buried within the hydrophobic core of the WIF domain, and the headgroup is accessible at the domain’s surface, in a manner reminiscent of ligand binding to lipocalins (Schiefner and Skerra, 2015). By contrast, the WIF domain of Drl-2 has a very well packed hydrophobic core (Figure 3C), in which no significant internal cavities could be found by the CASTp server (Tian et al., 2018). Indeed, the largest detectable cavity has a volume of just 77 Å^3^, compared with 1,510 Å^3^ for the cavity in WIF-1 that accommodates DPPC.

This key distinction between the WIF domains from Drl-2 and WIF-1 can be explained based on sequence and structural differences. On one side of the WIF domain β-sandwich (Figure 3D, left), the secondary structure elements of Drl-2 and WIF-1 overlay very well – including α1 plus strands β1, β2, β4, and β7, which are also among the most well conserved in sequence across WIF domains (Figure S3). The other half of the sandwich (Figure 3D, right) diverges much more. One key change is a 6-residue insertion between strand β2 and helix α2 in WIF-1 compared with Drl-2, colored magenta in Figures 3B, 3D, and S3. These additional residues form an extra 3_10_ helix in WIF-1, allowing this corner of the β-sandwich to be more splayed in WIF-1 than in Drl-2 (compare Figures 3A and B), and thus aiding accommodation of the DPPC molecule. Strand β3 in Drl-2 – which occludes the would-be DPPC binding site (Figure 3A) is not seen in the WIF-1 crystal structure because of this alteration. The second key change is in strand β9, which is interrupted in WIF-1 by a loop and 4-residue insertion (including a proline), shown in magenta in Figures 3B, 3D, and S3. Whereas the contiguous strand β9 of the Drl-2 WIF domain passes through the would-be DPPC binding site (Figures 3A,D) and occludes it, the insertion/bulge between β9 and β9’ in WIF-1 makes room for DPPC to bind to this WIF domain (Figures 3B,D). Thus, as a result of sequence differences between the class of WIF domains found in RYK/Drl family members and in WIF-1 orthologs respectively, the sDrl-2 structure described here shows that RYK/Drl WIF domains cannot accommodate a WNT-associated acyl chain when binding to their ligands, unlike the WIF domain from WIF-1 itself.

### WNT binding by ROR and RYK/Drl family extracellular regions

The structures of both the Nrk CRD and the Drl-2 WIF domain argue that the ECRs of these WNT-binding RTKs do not engage an attached acyl chain when they bind to their WNT ligands. The fatty acid molecule buried in the Nrk CRD core is unlikely to be capable of exchange with a WNT-attached fatty acid – contrasting with fatty acids bound in the readily accessible hydrophobic channels of FZD family CRDs (Figures S2B,C). Drl-2, as a representative of the RYK family, simply has no binding site to accommodate an acyl chain. Although these receptors have been shown to form complexes with WNTs (Reynaud et al., 2015; Ripp et al., 2018; Roy et al., 2018; Yoshikawa et al., 2003), it is important to ask whether the isolated ECRs are capable of interacting with the WNTs directly – for which we undertook ligand binding studies.

To investigate ligand binding by Nrk and Drl family proteins, we expressed and purified DWnt-5 (Eisenberg et al., 1992) in *Drosophila* Schneider-2 cells as described in Method Details. Note that DWnt-5 was also called DWnt-3 when it was first cloned (Fradkin et al., 1995; Russell et al., 1992). In our initial pull-down studies, DWnt-5 co-precipitated robustly with the histidine-tagged Drl ECR (sDrl_242_) or Drl WIF domain (sDrl_183_), as shown in Figure 4A. As a control, DWnt-5 did not co-precipitate significantly with the human PTK7 ECR. Pull-down experiments with the *Drosophila* ROR family ECRs gave substantially weaker signals than Drl, and significant levels of DWnt-5 were only seen in pull-downs of histidine-tagged s-dRor, consistent with a previous report (Ripp et al., 2018), but not sNrk (Figure 4A). In agreement with these findings, whereas Drl family proteins showed robust DWnt-5 binding in surface plasmon resonance (SPR) experiments as described below, we could not reliably detect strong DWnt-5 binding to sNrk using SPR. Turning to mammalian ROR family proteins for pull-down experiments, we produced murine WNT-5a in Expi293 cells as described previously (Speer et al., 2019), and found that the histidine-tagged ECR from human ROR2 could co-precipitate mWNT-5a (Figure 4B), in agreement with previous work (Billiard et al., 2005; Oishi et al., 2003) – although the ROR1 ECR could not. Importantly, mutating the acylated serine in mWNT-5a to alanine (S244A) did not prevent ROR2 from binding WNT-5a in this assay. Together with the binding of DWnt-5 to s-dRor1 in Figure 4A, this result supports the argument that WNT acylation is not crucial for WNT binding to ROR family CRDs.

**FIGURE 4.**
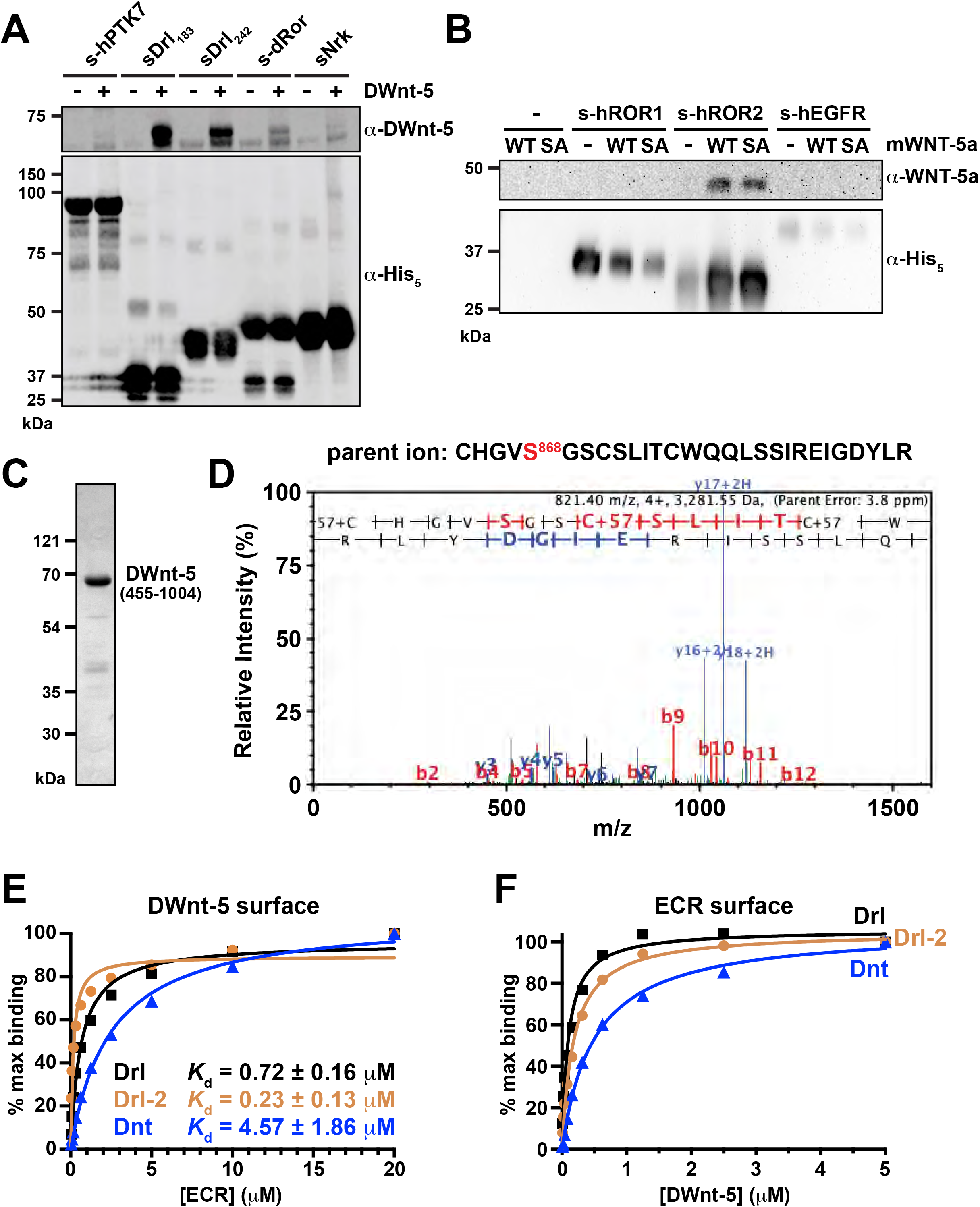
Binding of WNTs to ROR and RYK/Drl family ECRs. **(A)** Pull-down experiment as described in Method Details, showing that histidine-tagged sDrl_183_ and sDrl_242_ bind robustly to DWnt-5 in solution when precipitated with Ni-NTA beads and analyzed by immunoblotting with anti-DWnt-5. The dRor ECR (s-dRor) binds less robustly, and sNrk still more weakly. Lack of binding to the PTK7 ECR is shown as a control. Data are representative of at least 3 biological repeats. **(B)** Pull-down experiment showing that the histidine-tagged hROR2 ECR similarly precipitated mWNT-5a from conditioned medium from Expi293 cells expressing it. Neither the hROR1 ECR nor a human EGFR ECR control precipitated mWNT-5a in parallel experiments. Note the lack of effect of an acylation site (S244A) mutation in mWNT-5a, as discussed in the text. Data are representative of at least 3 biological repeats. **(C)** Coomassie-stained SDS-PAGE gel of purified DWnt-5, run after the size-exclusion chromatography step. **(D)** MS/MS product ion spectrum of 4+ charged ion at m/z 821.4, corresponding to the non-acylated peptide containing S868 of purified DWnt-5, which is the putative acylation site of this WNT ortholog. As described in the text, the ability to see this peptide at this abundance, alongside the monomeric behavior of DWnt-5, argues that it is not acylated. **(E)** SPR studies of Drl family ECRs binding to DWnt-5 immobilized on a sensorchip as describe in Method Details. Fit Kd values from at least 3 biological repeats are quoted ± standard deviation. **(F)** Corresponding reverse SPR experiment, with DWnt-5 (in solution) binding to immobilized sDrl, sDrl-2 and sDnt. Representative of at least three repeats. See also Figure S4.

### Recombinant DWnt-5 is not acylated, but binds tightly to Drl family WIF domains

Having detected robust binding of recombinant DWnt-5 to sDrl as described above, we purified the ligand to investigate the interaction in more detail and to assess the importance of DWnt-5 acylation. As shown in Figure S4A, DWnt-5 contains a long (∼550 aa) N-terminal ‘prodomain’ not seen in other WNT proteins, plus a ∼150 amino acid insert within its WNT homologous domain (Eisenberg et al., 1992; Russell et al., 1992). Early studies showed that DWnt-5 is proteolytically processed when expressed in an imaginal disc cell line to yield a predominant ∼80 kDa secreted species (Fradkin et al., 1995). DWnt-5 purified after expression in Schneider-2 (S2) cells was almost pure by Coomassie-stained SDS-PAGE (Figure 4C), and runs at just under 70 kDa (66 kDa by MALDI mass spectrometry). The protein was found by N-terminal sequencing to begin at residue 455 (sequence `SQPSIS`), which is ∼100 residues before the WNT homologous domain (Figure S4A). This protein, called DWnt-5(455-1004), is glycosylated at three sites that were identified by mass spectrometry as N484/485 (KVSME**NN**TSVTD), N724 (VDAK**N**DTSLV) and N952 (RVCHK**N**SSGLE). Deglycosylation with PNGase F reduces the apparent molecular weight of DWnt-5(455-1004) in SDS-PAGE to ∼64 kDa (Figure S4B), compared with a predicted value of 62 kDa. Importantly, mass spectrometry of a tryptic digest of DWnt-5(455-1004) clearly identified a peptide fragment containing unmodified S868, the putative lipid modification site of DWnt-5. The tryptic peptide extending from C864-R884 (Figure 4D) had a mass corresponding to the unmodified peptide (other than addition of iodoacetamide at each cysteine). The facts that this peptide was detected with similar abundance to other DWnt-5 peptides, and that the acylated derivative could not be detected, argue that recombinant DWnt-5(455-1004) produced in S2 cells is not lipid modified at S868. This lack of lipid modification is also consistent with the solubility of DWnt-5 without detergent, and its migration as a monomeric protein in size exclusion chromatography (see Method Details).

We used SPR to assess binding of DWnt-5(455-1004) to Drl family ECRs. The ECRs of Drl, Drl-2 and Dnt all gave robust binding signals in SPR studies, whether DWnt-5 was immobilized on the Biacore CM5 sensor chip and purified ECR was flowed across this surface (Figure 4E) or (conversely) the ECR was immobilized and DWnt-5 flowed across the surface (Figure 4F). Representative sensorgrams are shown in Figures S4C and D. The mean K_D_ for binding of the Drl ECR (sDrl_242_) to immobilized DWnt-5 across multiple repeats was 0.72 ± 0.16 μM. The WIF domain alone (sDrl_183_) bound with essentially the same affinity in parallel experiments (Figure S4E). The Drl-2 ECR bound DWnt-5 with similar affinity (0.23 ± 0.13 μM), and the Dnt ECR bound ∼10 fold more weakly. We further showed that this receptor binding was mediated by the Wnt homologous region of DWnt-5. As shown in Figure S4F, sDrl_1-242_ binding was unaffected when the large insert in the WNT homologous region (residues 681-838) was replaced with the corresponding 13-aa insert (RERSFKRGSREQG) seen in Wnt-5 from the ant *Harpegnathos saltator* (Bonasio et al., 2010) to generate DWnt-5_Δinsert_

These data show that the *Drosophila* RYK/Drl family receptors bind DWnt-5 with affinities typical for RTK ligand binding, despite both the absence of a lipid modification on DWnt-5, and the absence of an acyl-chain docking site in the ECR of the receptor.

### DWnt-5 binding does not promote sDrl dimerization

Since most RTKs undergo dimerization upon binding to their ligands (Lemmon and Schlessinger, 2010), we asked whether DWnt-5 might induce formation of sDrl dimers. Indeed, it has been reported that WNT-5a promotes hROR2 homodimerization (Liu et al., 2008) and that DWnt-5 drives Src64B recruitment to Drl by enhancing homodimerization of the receptor (Petrova et al., 2013). We used an in vitro pull-down assay to assess sDrl dimerization upon DWnt-5 binding (Figure 5A). Three different proteins were used: wild-type DWnt-5, FLAG-tagged sDrl_242_, and V5-tagged sDrl_242_. We incubated these proteins, either individually or in combination, with anti-FLAG conjugated to agarose beads and examined whether they could be pulled-down as a complex. DWnt-5 was efficiently pulled down by anti-FLAG when the FLAG-tagged sDrl_242_ was present, consistent with the data in Figure 4A, but the small amount of V5-tagged sDrl_242_ seen in anti-FLAG pull-downs did not change with addition of FLAG-tagged sDrl_242_ and/or DWnt-5 (compare lane 7 in lower blot of Figure 5A with lanes 3-5). Thus, these experiments argue that DWnt-5 does not induce sDrl dimerization – consistent with the previous suggestion that the transmembrane domain is likely to be the major driver of Drl dimerization (Petrova et al., 2013). We also note that the packing of sDrl-2 in crystals did not suggest any significant modes of ligand-independent ECR dimerization.

**FIGURE 5.**
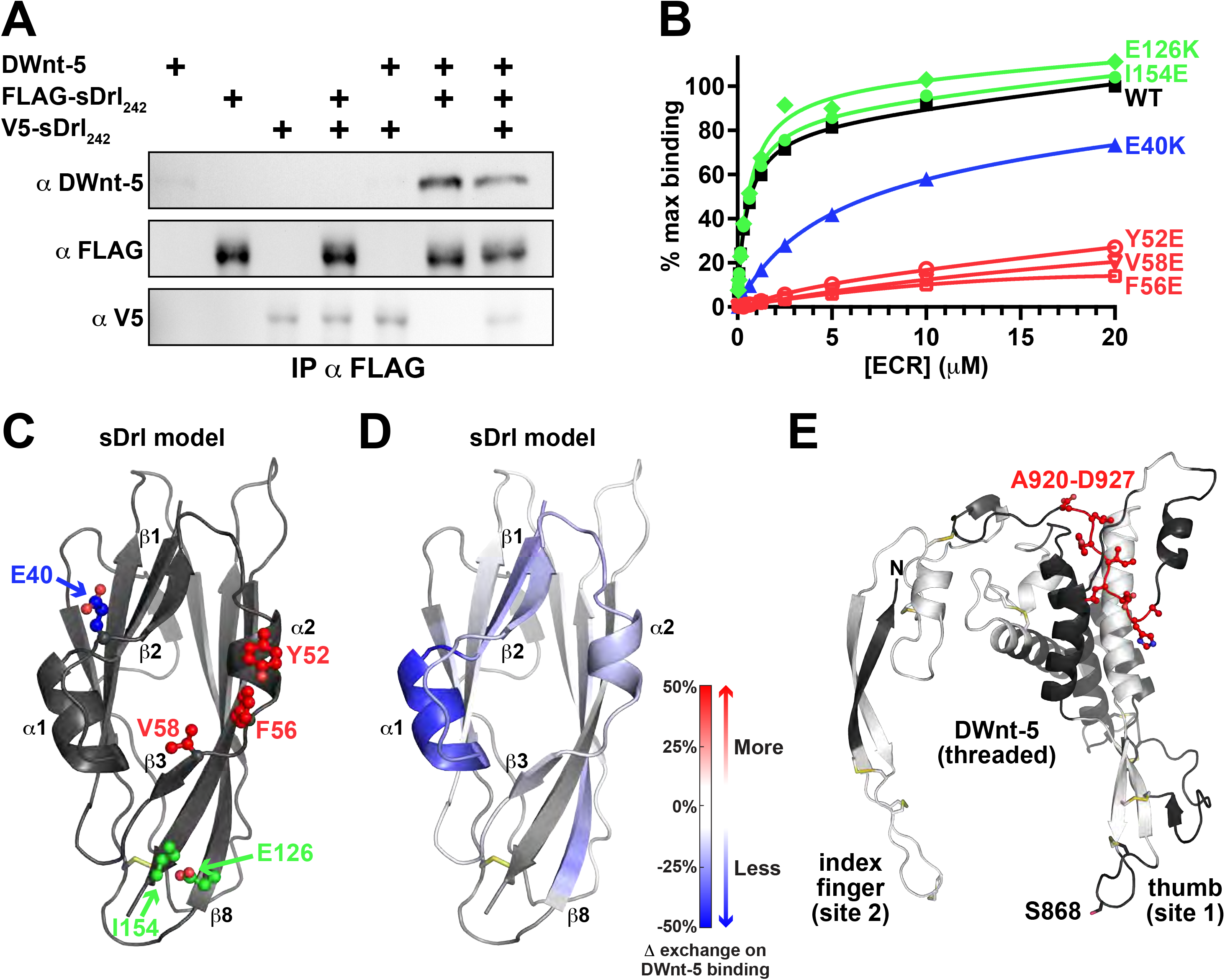
Locating the DWnt-5 binding site on Drl, and Drl binding site on DWnt-5. **(A)** Pull-down experiments indicate that DWnt-5 does not cause sDrl_242_ dimerization. FLAG-tagged and V5-tagged versions of sDrl_242_ were incubated with DWnt-5 as described in Method Details and immunoprecipitated with anti-FLAG. The quantity of V5-sDrl was not increased by the presence of FLAG-sDrl with- or without DWnt-5, arguing that they do not form dimers on WNT binding. **(B)** SPR data showing that Y52E, F56E, and V58E mutations (red) in sDrl_242_ greatly impair binding to immobilized DWnt-5. An E40K mutations (blue) has an intermediate effect, and E126K or I154E mutations have no detectable effects. Data represent at least 2 biological repeats. **(C)** Cartoon representation of an sDrl WIF domain model (based on the sDrl-2 WIF domain structure), showing the location of the mutations, color coded as in B. **(D)** Cartoon of sDrl model colored by the change in ‘weighted relative difference’ in HDX at the 1,000 s time point upon DWnt-5 binding, as described in Method Details. Regions in β1, α1, β2, α2 and to some extent β8 show some protection – supporting the location of the DWnt-5 binding site suggested by mutational analysis. **(E)** Limited HDX study of DWnt-5 changes upon binding sDrl in the same experiment. As described in the text, the large number of disulfides limit peptide coverage of the DWnt-5 protein. Grey/white areas are not seen in the recovered peptides. Only regions colored black were seen among the peptides (<50%), but showed no significant changes in HDX. Only the region colored red: A920-D927 in this threaded (xWnt8-based) model (Kelley et al., 2015) of DWnt-5 showed any evidence for reduced HDX. See also Figure S5.

### Location of the DWnt-5 binding site on RYK/Drl family WIF domains

We next took two parallel approaches to identify which surface of the Drl family WIF domains is responsible for the strong DWnt-5 binding seen by SPR. In the first, we mutated conserved residues in the Drl WIF domain and directly assessed their effects on DWnt-5 binding. In the second, we used hydrogen-deuterium exchange mass spectrometry (HDX-MS) to locate regions in the Drl WIF domain that are protected upon binding to DWnt-5.

Aligning the sequences of WIF domains from Drl, Drl-2 and Dnt with those from RYK/Drl family members of humans and model organisms identified several highly conserved surface residues marked with asterisks in Figure S3. These include Y52 and F56 of Drl (equivalent to Y57 and F61 in α2 of Drl-2), which are conserved in all RYK/Drl WIF domains, and E40, V58, E126, and I154 of Drl (equivalent to D45, V58, Q132, and I159 in Drl-2), which are all conserved in type. We individually mutated these six residues in sDrl_242_, and used SPR to assess the consequences for binding to immobilized DWnt-5. As shown in Figure 5B, glutamate substitutions at Y52, F56, or V58 (equivalent to Y57, F61, and V63 in Drl-2) essentially abolished DWnt-5 binding. Mutating E40 in sDrl to lysine (equivalent to D45 in Drl-2) had an intermediate effect, reducing affinity by ∼10-fold, and E126K or I154E mutations in sDrl (equivalent to Q132K and I159E in Drl-2) had no effect. Mutating L41 or Y42 in sDrl (L46 or F47 in Drl-2) caused protein aggregation, and so could not be studied. These mutagenesis studies implicate the surface that includes the α1/β2 loop, α2, and β3 in DWnt-5 binding (Figure 5C). Consistent with these results, HDX-MS analysis of sDrl showed that DWnt-5 binding to sDrl led to the greatest protection from backbone amide proton exchange in regions encompassing β1-α1-β2, α2, and β3 (Figure 5D; see also Figure S5A). Additional protection was also seen at the beginning of strand β8 – an area that has several well-conserved surface side-chains. With the exception of strand β8, these same parts of the Drl WIF domain showed the greatest backbone amine proton exchange in the absence of DWnt-5 (Figures S5B,C), suggesting that they are the most dynamic parts of the structure and are therefore likely alter conformation to accommodate DWnt-5 binding. The residues involved in DWnt-5 binding to Drl are in the same area of the WIF domain as those shown to be important for WNT3a binding to the WIF-1 WIF domain (Malinauskas et al., 2011) – in the region surrounding the splayed corner of the WIF domain β-sandwich that also allows accommodation of DPPC in WIF-1.

### Possible binding site for Drl on DWnt-5

We also attempted to gain insight into the binding epitope on DWnt-5 responsible for its binding to Drl family ECRs. Unfortunately, the 11 disulfide bonds, formed by 22 cysteines in the WNT homologous region of DWnt-5 limited MS peptide coverage quite substantially. HDX-MS analysis requires these disulfides to be reduced in order to assess mass changes in the individual peptic peptides, and it is very difficult to fully reduce the disulfides under the low temperature (and low pH) conditions required to minimize back exchange of amide protons after quenching of the reaction – even with very high levels of reducing agent (Bobst and Kaltashov, 2014). We tried several different approaches to optimize peptide coverage, but were not able to extend beyond ∼50% sequence coverage in the WNT homologous domain (Figure S6). None of the peptides that could be accurately assessed showed significant differences in exchange with- or without bound sDrl, potentially excluding these regions from being key parts of the receptor binding site. Notably, these included the peptide `HGVSGSCS` that covers residues 865-872 and includes the potential palmitoleoylation site, suggesting that this ‘thumb’ region (Figure 5E) may not be involved in Drl binding. Only one region showed any degree of protection, with peptides (in a non-disulfide bonded region) spanning residues 920-927 (`AHDLIYLD`) and beyond (Figure 5E). Although the poor coverage prevents us from being able to define the Drl-binding epitope on DWnt-5 with any certainty, these data suggest that it may differ from the region on WNTs that is engaged by FZD CRDs, possibly involving the opposite face of the molecule from the ‘palm’ described by Janda et al. (2012).

Intriguingly, the one region of DWnt-5 that can be implicated in Drl binding is in the ‘linker’ region between the large N-terminal domain (NTD) and smaller C-terminal domain (CTD) of the WNT molecule (Chu et al., 2013; Janda et al., 2012) – which has been shown to be required for co-receptor binding. Chu et al. (2013) showed that this region of WNT3a binds to LRP6. It was also recently reported that the WNT7-specific co-receptor Reck recognizes a similar region on WNT7a (Eubelen et al., 2018). Mutations in WNT7a that disrupt Reck binding were found to be concentrated in the linker region (aa 241-271 in hWNT7a) corresponding to the red-colored sequence in Figure 5E. These data suggest that RYK/Drl family receptors may recognize epitopes on WNTs distinct from those bound by FZDs, consistent with a co-receptor function.

### The Drl/DWnt-5 binding interface is required for Drl signaling in the *Drosophila* ventral nerve cord

Drl controls axon guidance in the developing central nervous system of *Drosophila* embryos (Callahan et al., 1995), specifically controlling which tracts axons use to cross the ventral midline. Drl is normally expressed in neurons that cross through the anterior commissure (AC), whereas those that cross through the posterior commissure (PC) normally do not express Drl, but do express DWnt-5. DWnt-5 in the PC functions as a repulsive signal, causing neurons that misexpress Drl to be redirected through the AC instead (Bonkowsky et al., 1999). This commissure switching by PC neurons that ectopically express Drl provides a useful quantitative in vivo assay for Drl function (Fradkin et al., 2004; Petrova et al., 2013; Yoshikawa et al., 2003), as summarized in Fig, 6A.

We ectopically expressed wild-type Drl or an F56E-mutated variant (which has lost the ability to bind DWnt-5) in Eg^+^ neurons that normally cross the ventral midline through the PC (left in Figure 6A), to assess whether they switch commissures. As shown in Figure 6B, expression of wild-type Drl resulted in robust commissure switching quantitated in Fig, 6C – whereas expression of the F56E variant did not (Figure 6B, right and Figure 6C). This finding indicates that mutating F56 impairs in vivo function of Drl as well as its in vitro binding to DWnt-5, and identifies F56E as a useful loss-of-function mutation for further in vivo dissection of RYK/Drl signaling.

**FIGURE 6.**
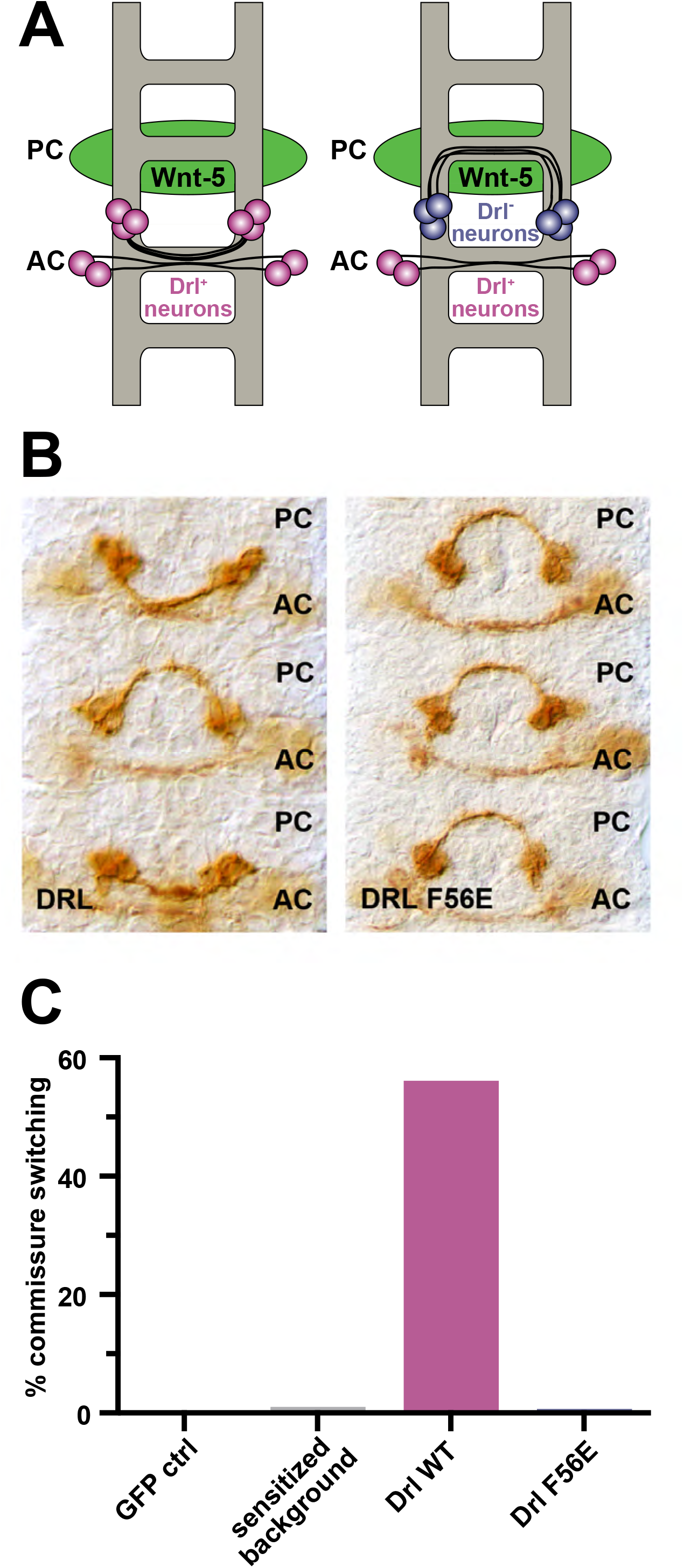
A DWnt-5 binding-deficient Drl variant abolishes commissure switching in vivo. **(A)** Schematic depiction of the Eg-Gal4 xUAS-Drl assay. One set of Eg^+^ neurons (blue) crosses the ventral midline via the posterior commissure (PC), and the other (magenta) through the anterior commissure (AC). DWnt-5 (green) is expressed predominantly by neurons that pass through the PC, which normally do not express Drl. When wild-type Drl is expressed ectopically in the Eg^+^ lineage, neurons that normally cross in the PC switch to cross in the AC, just below, to avoid the repulsive DWnt-5 signal. PC-to-AC switching of the Eg^+^ neurons therefore represents an assay for wild-type Drl function (Fradkin et al., 2004; Petrova et al., 2013; Yoshikawa et al., 2003). The penetrance of this phenotype is dependent on the levels of wild-type Drl expression; high levels result in essentially complete switching in all segments. We previously generated a Eg-GAL4/UAS-Drl line that serves as sensitized background. Individuals bearing single copies of the driver and the UAS-Drl insert display little commissure switching, whereas those with two copies display significantly increased levels of switching (Petrova et al., 2013). **(B)** Representative photographs of embryonic ventral nerve cords expressing wild-type (left) versus F56E-mutated (right) Drl in the single-copy sensitized background. Commissure switching occurred in wild-type Drl-expressing cases (2 of 3 commissures shown) but not in the F56E-expressing background. **(C)** Quantitation of commissure switching in controls, and embryos expressing wild-type Drl versus F56E-mutated Drl in the sensitized background. All animals, including the sensitized background control, contain single copies of the Eg-GAL4 transgene and a MYC-tagged wild-type DRL UAS transgene. The GFP control, wild-type, and F56E mutated transgenes are all present as a single copy. Expression of wild-type Drl results in robust commissure switching, whereas F56E-mutated Drl supports switching only at low background levels. At least 300 hemisegments were scored for each genotype. See also Figure S6.

## CONCLUSIONS

Although it seems clear that ROR and RYK family members of the RTK superfamily play important roles in WNT signaling, their transmembrane signaling mechanisms are not yet understood (Green et al., 2014; Roy et al., 2018; Stricker et al., 2017). The presence of FZD-like CRD and WIF domains in the ECRs of ROR and RYK family receptors respectively initially suggested that WNT binding to these RTKs might resemble that seen for FZDs (Hirai et al., 2019; Janda et al., 2012) and proposed for WIF-1 (Kerekes et al., 2015; Malinauskas et al., 2011). Our crystal structures of CRD and WIF domains from *Drosophila* ROR and RYK/Drl family members, however, argue that they have their own unique characteristics, particularly with respect to the role of WNT-associated acyl chains in receptor engagement. The absence of the acyl chain docking site on the Nrk CRD argues that this receptor does not engage WNTs (or their acyl chains) as described for FZDs. The absence of a lipid binding site in the Drl WIF domain – and its robust binding to non-acylated DWnt-5 – implicates different epitopes on the ligand in binding to this receptor (as do our HDX-MS studies).

Several studies indicate that ROR family members form WNT-dependent complexes with FZDs (Grumolato et al., 2010; Li et al., 2008; Nishita et al., 2010; Oishi et al., 2003). This would argue that the ROR and FZD CRDs recognize different regions of the WNT surface, allowing WNTs to crosslink FZDs and RORs – with the latter as co-receptors as suggested (Grumolato et al., 2010). Similarly, RYK appears to form WNT-dependent complexes with FZDs (Kim et al., 2008; Lu et al., 2004). Moreover, adding the CRD from secreted FZD-related protein-2 (sFRP2) in vitro efficiently blocks WNT binding to FZD5, but not to RYK (Schmitt et al., 2006). Early studies by Hsieh et al. (1999) also suggested that the WNT proteins interact differently with WIF-1 and FRPs, consistent with a distinct epitope on the WNT for WIF domain binding. By analogy with the WNT7a co-receptor Reck (Eubelen et al., 2018), and suggested by our HDX-MS data, this epitope might be in the linker region, close to the region implicated in LRP6 binding (Chu et al., 2013).

Much remains to be learned about WNT biology. Recent advances in understanding WNT/FZD interactions have opened up key approaches for defining the roles of different FZD subtypes when brought together with LRP5/6 as a common co-receptor in this complex signaling axis (Miao et al., 2020; Tao et al., 2019; Tsutsumi et al., 2020). Our results reveal the distinct structural characteristics of known WNT-binding modules in RTK ECRs, which we expect will provide an initial basis for designing approaches to understand how recruiting different co-receptors to a given FZD defines signaling function. Thus, these data represent an important step in dissecting the roles of different co-receptor complexes in WNT biology.

## STAR METHODS

Detailed methods are provided in the online version of this paper and include the following:

- **KEY RESOURCES TABLE**
- **CONTACT FOR REAGENT AND RESOURCE SHARING**
  - Materials Availability
  - Data and Code Availability
- **EXPERIMENTAL MODEL DETAILS**
  - Cell culture
  - *Drosophila melanogaster* studies
- **METHODS DETAILS**
  - Plasmid construction for recombinant protein expression
  - Protein production and purification
  - Crystallization and structure determination
  - in vitro pull-down assays for binding assessment
  - Surface Plasmon Resonance (SPR)
  - Small angle X-ray scattering (SAXS)
  - Mass spectrometry analysis for identification and post-translational modification
  - Hydrogen-Deuterium exchange-mass spectrometry (HDX-MS)
  - *Drosophila* commissure switching assays
- **QUANTIFICATION AND STATISTICAL ANALYSIS**
  - Structure determination and analysis
  - SPR data analysis
  - Analysis of HDX dynamics
  - Western blot image processing

## SUPPLEMENTAL INFORMATION

Supplemental information includes 6 figures and can be found in the online version of this article.

## ACKNOWLEDGEMENTS

We thank members of the Lemmon and Ferguson labs for comments on the manuscript, and Richard Gillilan (CHESS) and Kushol Gupta (UPenn) for help with SAXS data collection and analysis. This work was supported by NIGMS grants R35-GM122485 (to M.A.L.) and R01-GM031847 (to S.W.E), T32-GM007229 (to K.F.S.), an NSF Graduate Research Fellowship (DGE1122492 to J.B.S.) NSF grant MCB1020649 (to S.W.E.), and by the Deutsche Forschungsgemeinschaft (SO 1729/1-1 to A.S.). Synchrotron SAXS data were collected at MacCHESS beamline G1. CHESS was supported by NSF award DMR-0936384, and the MacCHESS resource by NIH/NIGMS award GM-103485. NE-CAT at the Advanced Photon Source (APS) is supported by a grant from NIGMS (P30 GM124165). APS is a U.S. Department of Energy (DOE) Office of Science User Facility operated for the DOE Office of Science by Argonne National Laboratory under Contract No. DE-AC02-06CH11357.

## AUTHOR CONTRIBUTIONS

F.S., J.M.M., L.G.F., and M.A.L designed the overall project. F.S., J.M.M., Z.W., and A.S. performed protein expression. F.S., J.M.M., K.H.P., and S.E.S. carried out crystallographic studies. J.M.M. and S.E.S. performed SAXS studies. Z.W. and F.S. performed HDX-MS – guided by S.W.E. and Z.-Y.K. – and J.B.S. assisted with analysis. F.S., J.M.M., A.S., and J.B.S. performed in vitro pull-down studies, and F.S., J.M.M., and K.F.S. undertook SPR experiments. L.G.F. and J.N.N. carried out in vivo experiments. M.A.L., F.S., and J.B.S. wrote the manuscript. All authors contributed to analysis of results and editing of the manuscript.

## SUPPLEMENTAL FIGURE LEGENDS

**FIGURE S1 – Related to Figure 1.**

**Structural features of ROR family ECRs**

(**A**) Most probable molecular envelope derived from SAXS analysis of the human ROR2 ECR as described in Method Details. The envelope is shown as a collection of spheres from the DAMAVER output, in two orthogonal views. Into the envelope have been docked the sNrk structure shown in Figure 1C (CRD plus Kr domain) plus an immunoglobulin (Ig) domain (cyan). This analysis shows that the Ig extends the ECR as a rod compared with sNrk.

(**B**). Pair distance distribution function or *P*(r) curve for s-hROR2 determined as described in Method Details. The maximum dimension (d_max_) of s-hROR2 is 135 Å, compared with an estimated 60-65 Å for sNrk and ∼45Å for the long axis of an Ig domain.

**(C)** Representative corrected scattering curve for s-hROR2 at 11.3 mg/ml (270 µM) in 25 mM MES, 150 mM NaCl, pH 6.0, with exposure time of 4 s, plotting intensity (*I*) plotted against *q* (4πsinθ/λ, where 2*q* is the scattering angle).

**(D)** Guinier analysis of the sample shown in **C**, with the Guinier region (*q*\**R*_g_ < 1.3) marked and residuals of the fit (lower points) shown.

**(E)** Overlay of the Nrk/dRor2 Kringle domain with those determined by X-ray crystallography for hROR1 (green; PDBID: 6BAN) and hROR2 (yellow; PDBID: 6OSH) in complex with potential therapeutic antibodies (Goydel et al., 2020; Qi et al., 2018). Two orthogonal views are shown, as marked.

**(F)** Illustration of the only dimer in the sNrk crystals that buries more than 400 Å^2^ of surface. Three orthogonal views are shown. One molecule is colored as in Figure 1C, and the other pink (CRD) and white (Kr). Water molecules are shown as red spheres. As can be seen in the top and bottom views, there is space between both the CRDs and the Kr domains in this dimer, with water between the molecules. Thus, there is no intimate dimer interface – consistent with the fact that sNrk showed no evidence for dimerization in size exclusion chromatography, SAXS, or analytical ultracentrifugation studies.

**FIGURE S2 – Related to Figure 2.**

**Sequence comparison of ROR family CRDs**

**(A)** Structure-based sequence alignment of *Drosophila* ROR family CRDs with those from human RORs, MuSK, and murine FZDs 4 and 8. Cysteines are shaded yellow/orange. Secondary structure elements are marked on the sequence for Nrk (purple) and mFZD8 (cyan), using the designation introduced by the Leahy lab (Dann et al., 2001). Residues circled in black contact the bound fatty acid in the Nrk, mFZD8 (Hirai et al., 2019; Janda et al., 2012) and mFZD4 CRD (DeBruine et al., 2017) structures. Those involved in fatty acid binding to the Nrk CRD that are conserved in other CRDs are colored green – noting that hROR1 and hROR2 conserve many of these residues. Basic residues in the α5/C-tail connector that ‘clamp’ the fatty acid headgroup in the Nrk CRD are conserved (albeit not precisely) in location in all other ROR CRDs (and MuSK) – and are colored blue – but are absent in FZD CRDs, which have a much shorter connection between the helix α4 and this region. This suggests that all ROR CRDs are capable of the fatty acid binding mode seen for Nrk. The α4 insert that breaks α4 (and creates α5 as described in the text) is marked with a red arrow.

**(B)** View of the Nrk CRD with transparent surface, with fatty acid-contacting residues colored red (side-chains shown as sticks) – showing that they form an internal binding site for the buried fatty acid.

**(C)** View of the mFZD8 CRD in the same orientation used for Nrk in **B**. A transparent surface is shown. The fatty acid-contacting residues, many of which are common to mFZD8 and Nrk are colored red (side-chains shown as sticks) and form a very clear hydrophobic ‘channel’ on the domain’s surface – in which the fatty acid resides.

**(D)** Cartoon representation of the hFZD5 CRD dimer structure (PDBID: 5URY), mediated by α1, α2, and α4 and a single fatty acid molecule that simultaneously lies in the hydrophobic channel of both CRDs (Nile et al., 2017).

**FIGURE S3 – Related to Figure 3.**

**Sequence comparison of WIF domains**

The WIF domains from *Drosophila* RYK/Drl family members are aligned with those of RYK orthologs from *Caenorhabditis elegans* (LIN-18), zebrafish (zRyk), *Xenopus laevis* (xRyk), mouse (mRYK), and human (hRYK), plus the human WIF-1 WIF domain. Alignments are guided by the structures of the sDrl-2 and WIF-1 WIF domains, for which secondary structure elements are shown at the top (in orange) and bottom (deep teal). Sequence conservation is colored from red (conserved in all 9 sequence) to blue (conserved in 5 of 9), revealing key areas of conservation. The inserted 3_10_ helix sequence (`MAPFTH`) adjacent to α2 is colored magenta, as is the proline-containing insert in β9 (`PQNA`). Glycosylation sites are circled in black, and residues mutated in sDrl for binding studies are marked with asterisks, colored according to whether they greatly disrupted binding (red), partly disrupted binding (blue), or had no effect (green).

**FIGURE S4 – Related to** Figure 4.

**Characteristics and interactions of DWnt-5**

**(A)** Schematic of the DWnt-5 protein. After the amino-terminal signal sequence (orange) is a long ‘prodomain’ that was found to be cleaved off when the protein is expressed in an imaginal disc cell line (Fradkin et al., 1995), yielding a mature secreted form of ∼80 kDa. As described in the text, the mature protein produced here in S2 cells begins at residue 455 (based on N-terminal sequencing). The protein also has a characteristic insert in its WNT homologous region, from 685 to 838, which does not appear to be involved in RTK binding.

**(B)** Coomassie-stained SDS-PAGE of purified DWnt-5 before and after treatment with PNGase F, showing that it is glycosylated, and the protein core is ∼64 kDa.

**(C)** Representative sensorgram from an SPR experiment in which 320 nM sDrl_242_ was injected on a sensorchip onto which DWnt-5 had been immobilized. Note that the signal comes back down to baseline after the injection, showing spontaneous dissociation of the sDrl protein.

**(D)** Representative sensorgram from the converse SPR experiment in which 160 nM DWnt-5 was injected on a sensorchip onto which sDrl_242_ had been immobilized. Note that the signal does not come back down to baseline after the injection in this case, necessitating the regeneration step mentioned in Method Details.

**(E)** SPR binding curves for binding of the complete sDrl ECR (sDrl_242_) and the WIF domain (sDrl_183_) to immobilized DWnt-5, showing no difference. Similarly, mutation of two glycosylation sites (necessary for protein behavior for HDX-MS analysis) had no influence on DWnt-5 binding. Data are representative of at least 2 biological replicates.

**(F)** SPR binding curves comparing binding of sDrl_242_ to immobilized mature DWnt-5 and DWnt-5 from which the ‘insert’ in **A** had been removed as described in Method Details (DWnt-5_Δinsert_). Loss of the insert has no detectable influence. Data are representative of at least 2 biological replicates.

**FIGURE S5 – Related to Figure 5.**

**HDX-MS analysis of sDrl binding to DWnt-5**

**(A)** Summary of HDX differences observed in sDrl_242_ in the presence of DWnt-5 protein at ∼25 μM as described in Method Details. The left-hand representation of the structure depicts coverage (black regions were detected), which was ∼75% of the WIF domain in HDX experiments. Missing peptides were in disulfide bonded regions. For the peptides seen, the percent exchange at the different time-points shown, as a result of DWnt-5 binding is colored according to the scale and mapped onto the structural model. Data for 1,000 seconds are shown in Figure 5D.

**(B)** Representation of HDX in unliganded sDrl_242_ across 5 different time points, showing that the regions surrounding the splayed corner of the sandwich (including α1 and α2) are among the most highly exchanging regions.

**(C)** Plot of exchange data for the 5 time points, with the x axis representing the median residue number of the peptide for which mean exchange is plotted as described (Sheetz et al., 2020).

**FIGURE S6 – Related to Figure 6.**

**Coverage of DWnt-5**

Peptide map showing the best coverage obtained for DWnt-5_Δinsert_ in HDX-MS studies, corresponding to 172 peptides (124 unique peptides). The positions of the 24 cysteines (all in disulfide bonds) in DWnt-5_Δinsert_ are marked at the bottom in gold text. Sequence numbers correspond to the DWnt-5 sequence in Uniprot (P28466). Note that coverage is very poor in the C-terminal part of the protein – across the entire C-terminal domain of the WNT protein, where there are 12 cysteines.

## KEY RESOURCES TABLE

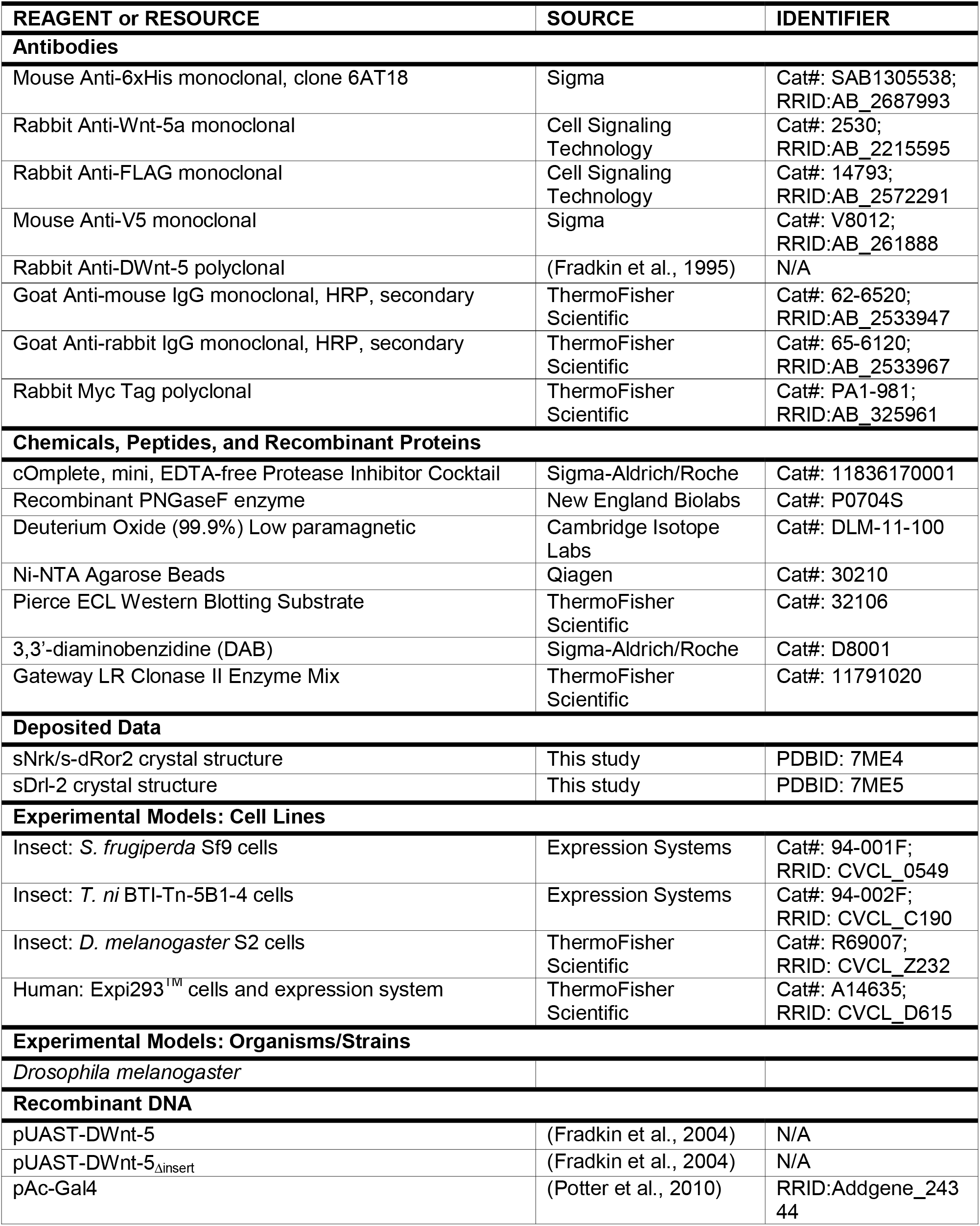

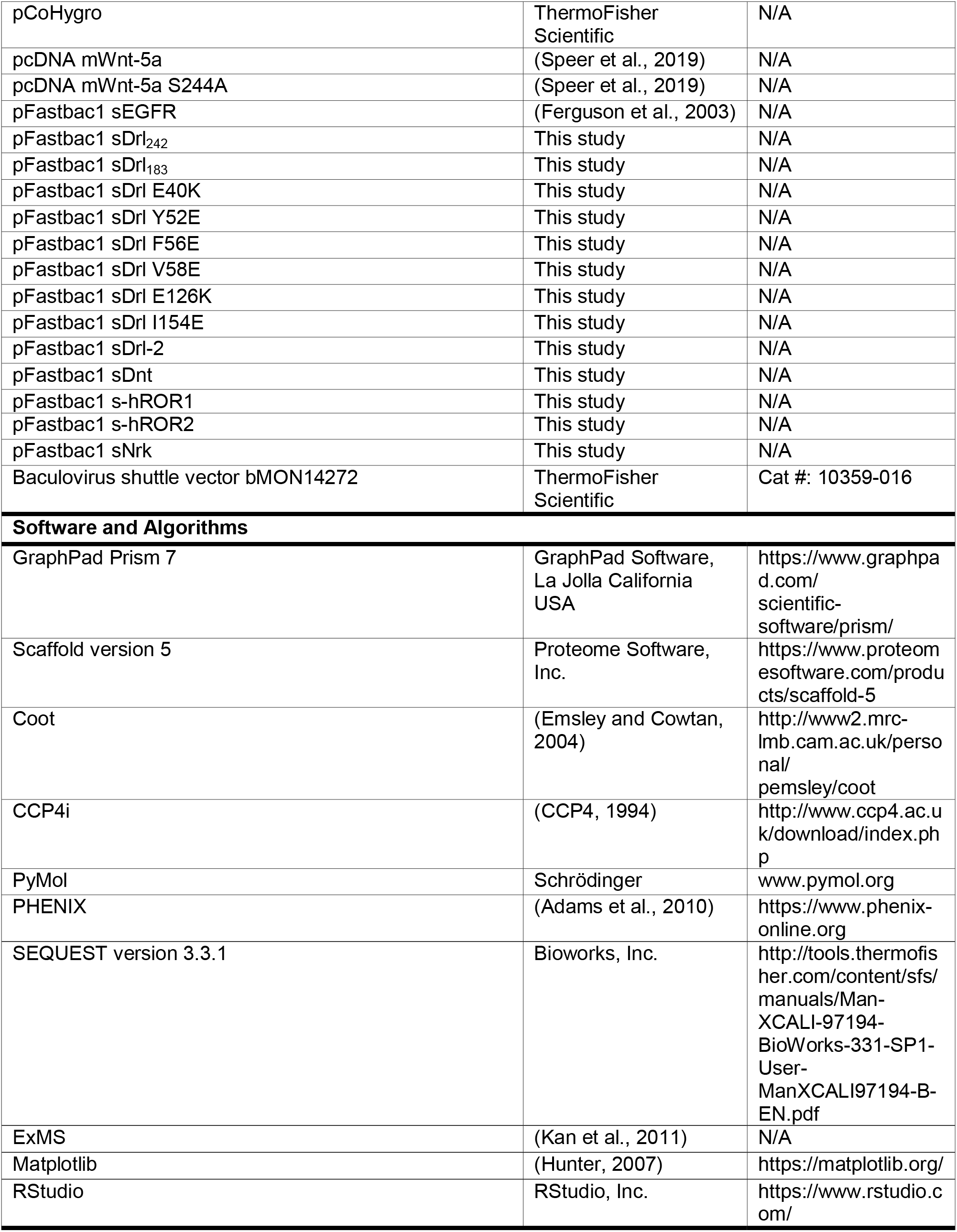

## STAR * METHODS

### CONTACT FOR REAGENT AND RESOURCE SHARING

Requests for further information or reagents may be directed to the Lead Contact, Mark A. Lemmon.

### Materials Availability

All unique and stable reagents generated in this study are available upon request.

### Data and Code Availability

PDB accession codes for the crystallographic coordinates and structure factors reported in this paper are: PDB: 7ME4 (Nrk/dRor2 ECR http://www.rcsb.org/structure/7ME4) and PDB: 7ME5 (sDrl-2 ECR http://www.rcsb.org/structure/7ME5). Original gel data have been deposited to Mendeley Data: https://doi.org/?

## EXPERIMENTAL MODEL DETAILS

### Cell culture

#### Insect cells

*Spodoptera frugiperda* Sf9 and *Trichoplusia ni* BTI-Tn-5B1-4 (High Five) cells were propagated at 27°C in serum-free ESF 921 Insect Cell Culture Medium (Expression Systems) containing 50 U/ml penicillin/streptomycin, and were used for production of secreted ECR proteins. Sf9 cells were originally established from immature ovaries of female *S. frugiperda* pupae, and BTI-Tn-5B1-4 cells from ovarian cells of the cabbage looper, *T. ni* and are also female. Schneider 2 (S2) cells were propagated at 24°C in Schneider’s Insect Medium (Sigma) supplemented with 10% fetal calf serum or in EX-CELL 420 serum-free medium (SAFC Biosciences). S2 cells were originally established from a primary culture of late stage male *D. melanogaster* embryos.

#### Mammalian cells

Expi293 cells (female) were grown in suspension in Expi293 Expression medium supplemented with 100 U/ml penicillin/streptomycin at 37°C and 8% CO_2_.

#### *Drosophila melanogaster* studies

Standard Drosophila husbandry practices were followed. Fly strains were maintained on standard molasses-cornmeal-yeast food and were kept at 25°C with a 12 h dark cycle. Embryos were collected from 0-24 h at 20-25°C, starting late in the afternoon onto grapefruit agar plates smeared with yeast paste subsequent to a 3 hour collection to purge females of retained older embryos. Crosses were established 24 hours prior to the start of embryo collection.

## METHOD DETAILS

### Plasmid construction for recombinant protein expression

cDNA fragments encoding receptor ECRs were subcloned into pFastBac1 for expression in Sf9 or *T. ni* cells. The coding regions corresponded to:

<u>s-dRor</u> (UniProtKB – Q24488), aa 1-313 – with C-terminal hexahistidine tag;

<u>sNrk/s-dRor2</u> (UniProtKB – Q9V6K3), aa 1-316 – with a C-terminal octahistidine tag;

<u>s-hROR1</u> (UniProtKB – Q01973), aa 1-406 – with C-terminal hexahistidine tag;

<u>s-hROR2</u> (UniProtKB – Q01973), aa 1-403 – with C-terminal hexahistidine tag;

<u>sDrl_183_</u> (UniProtKB – M9PDD9), aa 1-183 – with C-terminal hexahistidine tag;

<u>sDrl_242_</u> (UniProtKB – M9PDD9), aa 1-242 – with C-terminal hexahistidine tag;

<u>sDrl-2</u> (UniProtKB – Q7JQT0), aa 1-183 – with a spacer peptide (RPLESRGPFEGKPIPNPLLGLDSTRTG) followed by a C-terminal hexahistidine tag;

<u>sDrl-2/Xa</u> (UniProtKB – Q7JQT0), aa 1-183 – followed by a C-terminal Factor Xa (FXa)

cleavage site (`IEGR`), spacer peptide (ASGPFEGKPIPNPLLGLDSTRTG) and a hexahistidine tag;

<u>sDnt</u> (UniProtKB – M9PG69), aa 1-208 – with a spacer peptide (RPLESRGPFEGKPIPNPLLGLDSTRTG) followed by a C-terminal hexahistidine tag.

For expression of wild-type DWnt-5 protein (UniProtKB – P28455), we used a pUAST plasmid containing the open reading frame of wild-type DWnt-5. To generate DWnt-5_Δinsert_ protein, this plasmid was altered using site-directed mutagenesis to replace residues 681 to 838 of DWnt-5 was replaced with the short peptide sequence (RERSFKRGSREQG) found in the corresponding region of Wnt-5 from *Harpegnathos saltator* (Bonasio et al., 2010). Expression constructs for mWnt-5a expression were generated as described by Speer et al. (2019).

### Protein production and purification

#### sNrk for crystallization

Recombinant baculovirus was generated using the Bac-to-Bac system (ThermoFisher Scientific), and sNrk was expressed in Sf9 cells grown in ESF921 medium (Expression Systems). Conditioned medium was collected 3 days after infection, diafiltered against 20 mM NaKPO_4_, pH 8.0, containing 200 mM NaCl using a TFF2 10k-cutoff cartridge (Millipore), and loaded on to a Ni-NTA (Ni^2+^-nitrilotriacetate) column (Qiagen). The Ni-NTA resin was serially washed with the same buffer containing 10 mM, 20 mM and 30 mM imidazole (2 column volumes each), and then eluted in 20 mM NaKPO_4_, pH 8.0, containing 200 mM NaCl and 200 mM imidazole. After dialysis in this buffer to remove imidazole, protein was then loaded on to Fractogel EMD SO3^-^ cation exchange column (Millipore) in 20 mM MES, pH 6.0. After a step to 200 mM NaCl, protein was eluted with a gradient from 200 mM to 450 mM NaCl – eluting at 350-400 mM. Peak fractions were concentrated in an Amicon 10 concentrator, dialyzed into 20 mM HEPES, pH 8.0 containing 2.5 mM NaKPO_4_ plus 125 mM NaCl, and passed through a Bio-Rad Bio-Scale CHT2-I ceramic hydroxyapatite column. The flow-through was then concentrated and subjected to size exclusion chromatography on a Superose 6 column (GE Healthcare) in 20 mM HEPES, pH 7.5, containing 150 mM NaCl.

#### Other ROR family ECRs

s-dRor, s-hROR1 and s-hROR2 for pull-down experiments and SAXS experiments (s-hROR2 in Figure S1) were produced using essentially the same approach as described for sNrk, but omitting the hydroxyapatite chromatography step.

#### sDrl, sDrl-2, and sDnt proteins

Recombinant baculoviruses encoding Drl family ECRs were used to infect Sf9 cells (*T. ni* cells for sDrl-2 for crystallization) grown in ESF921 medium (Expression Systems). Conditioned medium was harvested three days after infection and subjected to extensive dialysis at 4°C against 20 mM HEPES pH 7.5, 150 mM NaCl. Proteins were then loaded onto Ni-NTA beads, which were washed twice with low imidazole buffer (20 mM HEPES pH 7.5, 150 mM NaCl and 15 mM imidazole) prior to elution of protein with buffer containing 20 mM HEPES pH 7.5, 150 mM NaCl and 100 mM imidazole. Drl family ECRs were further purified using an UnoQ anion exchange column (Bio-Rad), loading in 20 mM HEPES pH 7.5 containing 70 mM NaCl and using an elution gradient of 70 mM-1 M NaCl. Peak fractions were then subjected to size exclusion chromatography using a Superose 12 column (GE Healthcare) in 10 mM HEPES pH 7.5, containing 150mM NaCl. sDrl-2 for crystallization was subjected to Factor Xa protease (10 µg protease per mg of sDrl-2 in 1 ml for 1 h at room temperature) to remove the hexahistidine tag, and was then subjected to anion exchange and sizing. sDrl-2 for crystallization was also partially deglycosylated with PNGase F (New England BioLabs: 2,000 unit/mg sDrl-2) for 3 h at room temperature before sizing.

#### DWnt-5 purification

S2 cells were transfected with a mixture of three plasmids (*i*) pUAST-DWnt-5, (*ii*) pAc-Gal4 and (*iii*) pCoHygro (10μg:10μg:1μg) using the calcium phosphate method, and were selected in Schneider’s Insect Medium (Sigma) supplemented with 10% fetal calf serum (Sigma, Cat# F0643) and 300 μg/ml hygromycin (Cellgro) for 3 weeks. The Schneider’s Insect Medium was then replaced with EX-CELL 420 serum-free medium (SAFC Biosciences) for subsequent cell culture, and constitutive secretion of DWnt-5 into the medium was verified using DWnt-5 specific antibodies. For DWnt-5 expression, cells were seeded at 4×10^6^ cells/ml in spinner flasks. After 5 days of growth, medium (∼3 liters) was harvested and flowed through a 4 ml Fractogel SO3^-^ (EMD Millipore) cation exchange column at 4°C. The column was then washed twice with 10 ml of wash buffer (20 mM HEPES, pH 7.5 containing 250 mM NaCl), and DWnt-5 was eluted in 20 mM HEPES, pH 7.5 containing 900 mM NaCl in three washes of 4 ml each. The eluted protein was then diluted with 3 volumes of 20 mM HEPES, pH 7.5 to lower [NaCl] to <250 mM, and was loaded onto a second 2 ml Fractogel SO3^-^ AKTA column at room temperature, pre-equilibrated in 20 mM HEPES, pH 7.5 and 150 mM NaCl. DWnt-5 was eluted with a gradient from 150 mM to 1 M NaCl in this buffer, eluting at around 650 mM NaCl. The eluted fractions were then diluted again with 3 volumes of 20 mM HEPES, pH 7.5 and loaded onto a 2 ml CHT2-I hydroxyapatite column (Bio-Rad) equilibrated in 20 mM HEPES, pH 7.5 containing 150 mM NaCl, 2.5 mM NaH_2_PO_4_, and 2.5 mM K_2_HPO_4_. A gradient from 0-100% of Buffer B (20 mM HEPES, pH 7.5 containing 150 mM NaCl, 250 mM NaH_2_PO_4_, and 250 mM K_2_HPO_4_) was then applied. Eluted fractions were pooled, concentrated in a centrifugal concentrator, and subjected to size exclusion on a Superose 6 column (GE Healthcare) in 10 mM HEPES, pH 7.5 containing 150 mM NaCl. Then purified DWnt-5 protein could be flash frozen following addition of 10% glycerol with no significant aggregation or loss of sDrl-binding activity upon thawing.

#### mWnt-5a production

mWnt-5a-containing conditioned medium was prepared as described (Speer et al., 2019) by expressing wild-type or acylation-site-mutated mWnt-5a using the Expi293^TM^ Expression System (ThermoFisher Scientific) according to the manufacturers’ instructions. Expi293 cells were transfected with plasmids encoding wild-type or mutated mWnt-5a under control of a CMV promoter. The culture medium was harvested 96 h post-transfection and cleared by centrifugation for use in co-precipitation assays.

### Crystallization and structure determination

#### sNrk

For sNrk, the protein was concentrated to ∼3 mg/ml. Crystals were grown at 21°C using the hanging drop vapor diffusion method, mixing equal volumes of protein solution and well solution containing 50 mM Bis-Tris propane (pH 5.0), 20% PEG 3350. Crystals grew in a few days, and were cryoprotected by weaning into the same solution containing 18% sucrose. Frozen crystals diffracted to 1.75 Å at APS beamline 24-ID-E. Initial phasing was obtained using mr_rosetta (Terwilliger et al., 2012), which identified a starting model based on the MuSK CRD (Stiegler et al., 2009) and the 7^th^ Kr domain in apolipoprotein-a (Ye et al., 2001), with PDBIDs 3HKL and 1I71 respectively. Structural refinement and model building were then carried out iteratively using Refmac (CCP4, 1994), Phenix (Adams et al., 2010), and Coot (Emsley and Cowtan, 2004).

#### sDrl-2

Purified sDrl-2 protein was concentrated to >12 mg/ml for crystallization. Crystals were grown at 21°C using the hanging drop vapor diffusion method by mixing 1 µl protein solution with 1 µl well solution containing 100 mM Tris pH 8.5, 25% PEG 6000, 100 mM sodium acetate, and 15% glycerol. Crystals formed within 2 days and were frozen directly in liquid nitrogen. Data were collected to 1.95 Å resolution on a HighFlux HomeLab X-ray diffraction unit (Rigaku) with a Saturn 944 CCD detector, and were processed using HKL2000 software. Initial phasing was obtained by molecular replacement using a truncated poly-alanine model of the NMR structure (Liepinsh et al., 2006) of the WIF domain from hWIF-1 (PDB: 2D3J) as the search model in Phaser (CCP4, 1994). Structural refinement and model building were carried out using Refmac (CCP4, 1994), Phenix (Adams et al., 2010), and Coot (Emsley and Cowtan, 2004).

### in vitro pull-down assays for binding assessment

#### DWnt-5 interactions

Purified histidine-tagged sDrl, sNrk, or s-dRor at ∼0.5 μM (or the human PTK7 ECR as control) were mixed with a similar concentration of DWnt-5 and incubated with nutation for 30 min at 4°C in 20 mM HEPES pH 7.5, 150 mM NaCl. Ni-NTA beads were then added, pelleted, and washed extensively in buffer prior to immunoblotting with anti-DWnt-5 (upper panel of Figure 4A) or anti 6xHis (lower panel of Figure 4A).

#### hROR-mWnt-5a binding

Conditioned medium from Expi293 cells expressing wild-type or S244A-mutated mWnt-5a was added to ∼0.5 μM histidine-tagged s-hROR1 or s-hROR2 (or s-hEGFR_501_ as a control) in 20 mM HEPES pH 7.5, 100 mM NaCl for 30 min at 4°C. Ni-NTA beads were then added, pelleted, and washed extensively in buffer prior to immunoblotting with anti-Wnt5a (upper panel of Figure 4B) or anti 6xHis (lower panel of Figure 4B).

#### Assessment of DWnt-5-induced sDrl dimerization

Two variants of sDrl_242_ were generated, with a V5-tag (GKPIPNPLLGLDSTGHHHHHH) and FLAG-tag (DYKDDDDKGHHHHHH) respectively after residue 242, and were produced in Sf9 cells using the approaches outlined above. In a total volume of 400 μl, 200 nM of sDrl-V5 and 100 nM sDrl-FLAG were mixed with 300 nM DWnt-5 and 15 μg anti-FLAG M2 in 20 mM HEPES pH 7.5, 150 mM NaCl and 0.2% (w/v) BSA. After 30 min at 4°C, 150 μl of Protein G Dynabeads (Invitrogen) were washed and added to the mixture, which was incubated for an additional hour. Supernatant and Dynabeads were then separated using a magnet, and Dynabeads were resuspended in equal volumes of buffer for immunoblotting anti-DWnt-5 (upper panel in Figure 5A), anti-FLAG (middle panel in Figure 5A) and mouse anti-V5 (lower panel in Fig, 5A).

### Surface Plasmon Resonance (SPR)

SPR experiments were performed using a Biacore3000 instrument (GE Healthcare). DWnt-5 protein was immobilized on CM5 sensorchips using the amine coupling method recommended by the manufacturers, typically immobilizing ∼10,000 resonance units (RUs) onto the surface. Purified sDrl at a series of concentrations (4 nM – 20 μM; starting at the lowest concentration) in 10 mM HEPES (pH 8.0), 150 mM NaCl, 3 mM EDTA and 0.005% Surfactant P-20 was then injected at 10 μl/min at room temperature until steady state was reached. Following each injection, bound sDrl was allowed to spontaneously dissociate from the sensorchip surface in the same buffer. Steady-state signals were background-corrected by subtracting the signal obtained with a control surface. For estimation of binding affinities, SPR signal values were plotted against [sDrl] and fit to a simple single-site saturation-binding model in Prism 9.

For the reverse experiment (immobilizing sDrl homologues and flowing DWnt-5 protein across the resulting surfaces), purified ECRs at 25 µM (in 8 mM HEPES pH 7.5, 120 mM NaCl, 10% glycerol) were diluted 1:4 in 10 mM sodium acetate, pH 4.5, and flowed across an activated CM5 sensorchip surface for 9 min at 10 µl/min prior to quenching with 1 M ethanolamine. Approximately 10,000-14,000 RUs of each sDrl protein were thus immobilized. Purified DWnt-5 was then injected at a range of concentrations (10 nM – 5 µM) until steady state was reached (typically ∼7 min at 10 µl/min). In this case, regeneration with a 25 μl injection of 10 mM sodium acetate, pH 4.5 containing 500 mM NaCl was required between injections.

### Small angle X-ray scattering (SAXS)

The SAXS data used for Figures S1A-D were recorded using beam line G1 at CHESS, for s-hROR2 at 11.3 mg/ml (270 µM) in 25 mM MES, 150 mM NaCl, pH 6.0. Data were collected in 2010 on a custom 1024 x 1024 (69.78 μm) pixel CCD detector constructed by the Grüner group (Cornell University, Ithaca, NY, USA). Two-dimensional images were integrated using Data Squeeze 2.07 (Datasqueeze Software, Wayne, PA, USA) to give one-dimensional intensity profiles as a function of *q* (*q* = 4πsinθ/λ, where 2θ is the scattering angle). Measurements were taken at room temperature with a sample-to-detector distance of 1,175 mm. With a calibrated wavelength of 1.256 Å, scattering profiles covered a *q* range from 0.008 to 0.294 Å^-1^. The incident X-ray beam was collimated to a spot size measuring 0.5 x 0.5 mm^2^, which was significantly smaller than the opening of the sample cells. Exposure times ranged from 4 s with no attenuation, and measurements were made in triplicate unless otherwise noted. Capillary quartz sample cells holding approximately 35 μl were used, with volume oscillation during data collection to help protect from radiation damage.

Data were corrected for incident radiation and scattering from a buffer match (against which the sample had been dialyzed) to yield the scattering profile (Figure S1C) in which intensity (*I*) is plotted as a function of *q*. Guinier analysis (Figure S1D) was performed using PRIMUS (Konarev et al., 2003). Pair-distance distribution functions (Figure S1B) were generated from the scattering profiles using the program GNOM (Semenyuk and Svergun, 1991), and results corroborated using an automated implementation of the program called AUTOGNOM (Petoukhov et al., 2007). The maximum diameter of the protein (*d*_max_) was adjusted in 10 Å increments in GNOM to maximize the goodness-of-fit parameter. This analysis also yielded an *R*_g_ determination. Low resolution shapes/most probable envelopes were determined from SAXS data using the program DAMMIF (Franke and Svergun, 2009). Ten independent calculations were performed for each data set, using default parameters with no symmetry assumptions. The models resulting from these independent runs were superimposed using the program SUPCOMB based on the normalized spatial discrepancy (NSD) criterion – with NSD values of 0.6-0.7 (Kozin and Svergun, 2001). The ten independent reconstructions were then averaged and filtered to a final consensus model using the DAMAVER suite of programs (Volkov and Svergun, 2003).

### Mass spectrometry analysis for identification and post-translational modification

Mass spectrometry analysis of trypsinized DWnt-5 and other proteins – as well as small molecule analysis – was provided by the Proteomics Core Facility at the Children’s Hospital of Philadelphia, using standard protocols for gel purified protein identification. Protein was treated with 10 mM iodoacetamide to reduced disulfide bonds and alkylate free cysteines. For identification of glycosylation sites, PNGase F treatment in ^18^O water was performed as described (Cao et al., 2018) prior to LC MS/MS analysis.

### Hydrogen-Deuterium exchange-mass spectrometry (HDX-MS)

For HDX-MS analysis of sDrl, the exchange reaction was initiated by mixing the sDrl N63Q/N143Q double mutant protein stock (28.5 μM, in 20 mM HEPES pH 7.4, 200 mM NaCl) into a 96% D_2_O solution containing 150 mM NaCl at a ratio of 1:4 (v:v). For the sDrl/DWnt-5 complex, a protein mixture containing 28.5 μM sDrl and 29.5 μM DWnt-5 was similarly diluted into 96% D_2_O (150 mM NaCl). Final concentrations of sDrl and DWnt-5 in the exchange reactions were thus 5.7 μM and 5.9 μM, respectively. The pD of each HDX reaction solution was estimated to be 7.2 (pH_read_ + 0.4). HDX reactions were carried out at 0°C. At each time point (10 s, 10^2^ s, 10^3^ s, 10^4^ s and 10^5^ s), a 15 μl aliquot of the reaction mixture was quenched by adding 45 μl of quench buffer (1.5 M guanidine hydrochloride, 50 mM tris-(2-carboxyethyl)phosphine (TCEP), 0.8% formic acid and 10% glycerol). As controls, non-deuterated (‘all-H’) and fully deuterated (‘all-D’) samples were prepared in the same way. All samples were frozen in liquid nitrogen immediately after adding quench buffer. For experiments seeking DWnt-5 peptides, the quench buffer was switched to 500 mM TCEP, pH 2.8 with 500 mM glycine/HCl as buffer agent - to maximize the number of reduced peptides while not having significant back exchange effects.

Prior to data collection, frozen samples were quickly thawed on ice and injected at a flow rate of 100 μl/min into a thermoelectrically cooled chamber (Mayne et al., 2011). The sample was digested on an immobilized pepsin column within the cooled chamber. Digested peptides were flowed through a Piccolo C18 tap column (Higgins Analytical) to desalt the peptide fragments. An acetonitrile gradient (10-55% acetonitrile, 0.1% TFA) was then used to elute peptides from the trap column and into an analytical C18 column (5 cm x 0.3 mm, Targa 3 μm C18 resin, Higgins Analytical). The effluent was flowed directly to Thermo LTQ Orbitrap XL mass spectrometer for electrospray ionization. A tandem MS (CID mode) run was carried out for the ‘all-H’ sample in order to identify the sequences of digested peptides. SEQUEST (Bioworks, version 3.3.1) was used to identify peptides from the tandem MS data. The MATLAB-based data analysis tool ExMS (Kan et al., 2011) was used to validate peptide assignments and subsequently to compute the centroid of isotopic distribution of each deuterated peptide. The ‘All-D’ sample was included to calibrate back-exchange of the deuterated samples. Details of ExMS-based data collection and the data processing workflow are described by Kan et al. (2011). NumPy and Matplotlib (Hunter, 2007) were used to export the ExMS results to the Python environment for further analysis and plotting. To assess the differences in HDX rates within sDrl with- and without bound DWnt-5, the difference in number of exchanged deuterons was calculated for each peptide at each time point.

The maximum difference among all of the time points was further divided by the number of amide hydrogen atoms in the peptide to give a weighted relative HDX difference for each peptide, which was visualized in Figures 5D,E and S5 by color coding the sDrl homology model according to this ‘weighted relative difference’.

### *Drosophila* commissure switching assays

The F56E mutation was introduced into the Drl coding region in the pENTR vector using oligonucleotide-mediated mutagenesis, and the open reading frame was subsequently transferred into pTWM-attB (L.G.F., unpublished) using LR Clonase (ThermoFisher Scientific) to generate Drl-F56E-(6x)-Myc. Transgenic lines of both wild-type Drl and Drl-F56E were then generated by phC31-mediated transgenesis using the pBac{yellow[+]-attP-9A}VK00027 stock (attP inserted at cytogenetic location 89E11) at Bestgene, Inc. to ensure equivalent expression of both species. A representative line of each was then crossed with the eg-GAL4 driver line with an insert of wild-type pTWM-Drl that shows minimal posterior to anterior commissure switching of the Eg+ neurons on its own (“sensitized background”). Embryos were collected from 0-24 hours at 20-25°C, starting late in the afternoon onto grapefruit agar plates smeared with yeast paste subsequent to a 3 h collection to purge females of retained older embryos. Crosses were established 24 hours prior to the start of embryo collection. 0-24 h embryos were collected, devitellinized and stained with rabbit anti-Myc (ThermoFisher Scientific) followed by HRP-conjugated goat anti-rabbit (ThermoFisher Scientific) and were visualization by incubation with a 3,3’-diaminobenzidine (DAB)/hydrogen peroxidase solution. Embryos were cleared by stepwise incubation with increasing concentrations of glycerol in phosphate-buffered saline, ventral nerve cords were dissected and mounted on slides and were then scored blinded to genotype. Controls included the sensitized background stock and pTWM-GFP (inserted at the same attP site) in the sensitized background.

## QUANTIFICATION AND STATISTICAL ANALYSIS

### Structure determination and analysis

Statistics for the structural models are provided in Table 1. Analysis of molecular contacts and RMSD values were calculated using the CCP4 software package (CCP4, 1994).

### SPR data analysis

Where *K*_d_ values are quoted, experiments were performed at least three times with different protein preparations – to achieve at least 3 biological replicates. *K*_d_ values are quoted ± SD.

### Analysis of HDX dynamics

Raw mass spectra of undeuterated controls were used for peptide identification using SEQUEST (Bioworks, Version 3.3.1). All raw spectra for each peptide, labeling condition, drug condition, and charge state were then manually assessed for quality and for accurate peak assignment, at which point poor quality or incorrectly assigned peaks were unassigned. The MATLAB-based software ExMS (Kan et al., 2011) was used to validate peptide assignment and to determine average mass shifts of centroids and their standard deviations to calculate percent uptake for each time point relative to a fully deuterated standard as described in Method Details.

### Western blot image processing

Raw images from a Kodak Image Station (Figures 4B,C, 5A, and S4B) or LI-COR Odyssey Fc imager (Figure 4A) were imported in Adobe Photoshop, and the ‘Levels’ function used to apply a linear correction (bring up background, bring down upper limit) so that the darkest points of all images are black, and the background is brought into the visible grey scale in order to register all features in the image.

## DATA AND SOFTWARE AVAILABILITY

Accession numbers for the coordinates and structure factors for the sNrk/s-dRor2 and sDrl-2 structures reported in this paper are PDB: 7ME4 and 7ME5, respectively.

