## Supplementary figures and images for "ROR and RYK extracellular region structures suggest that receptor tyrosine kinases have distinct WNT-recognition modes"

### Supplementary Figure 1

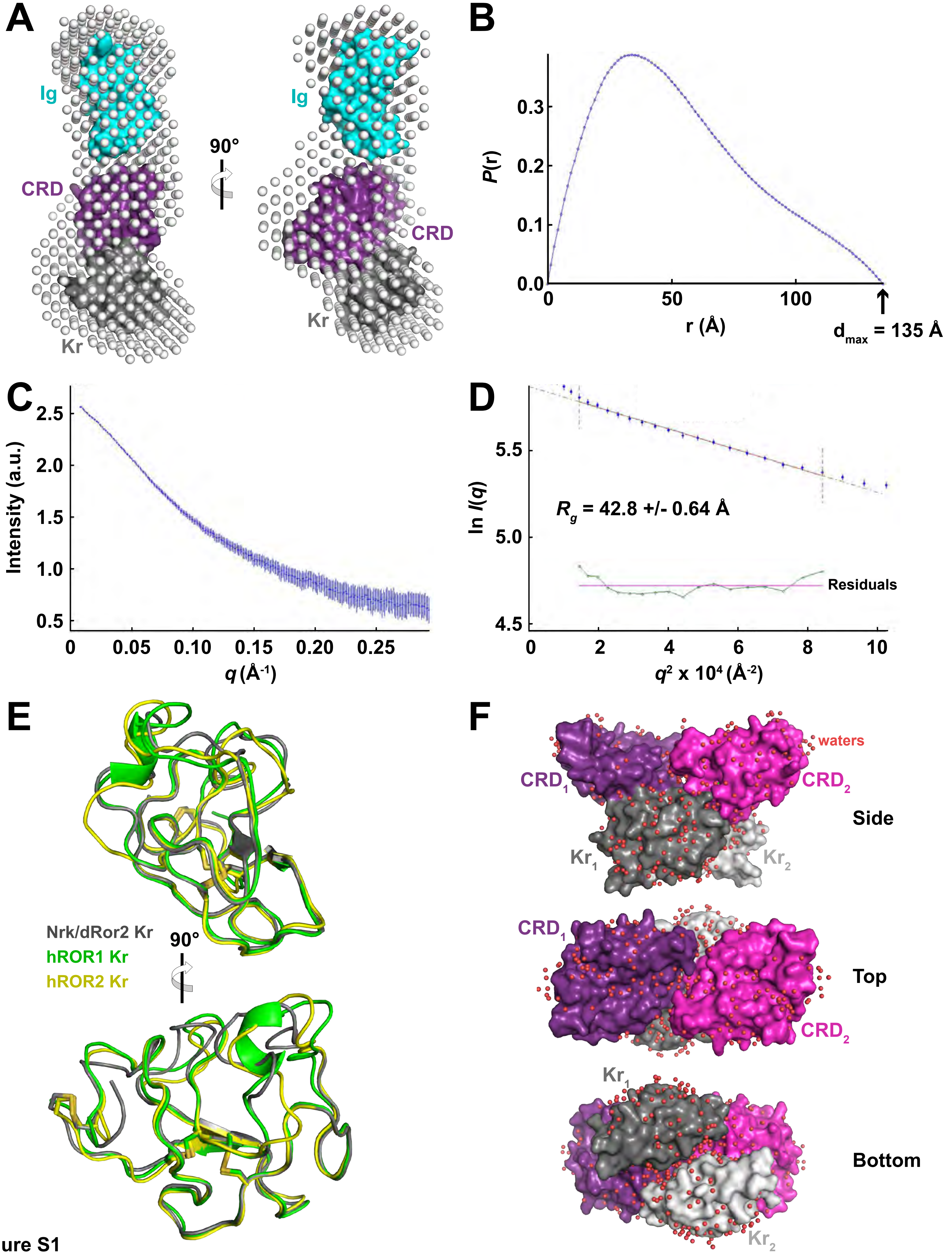

### Supplementary Figure 2

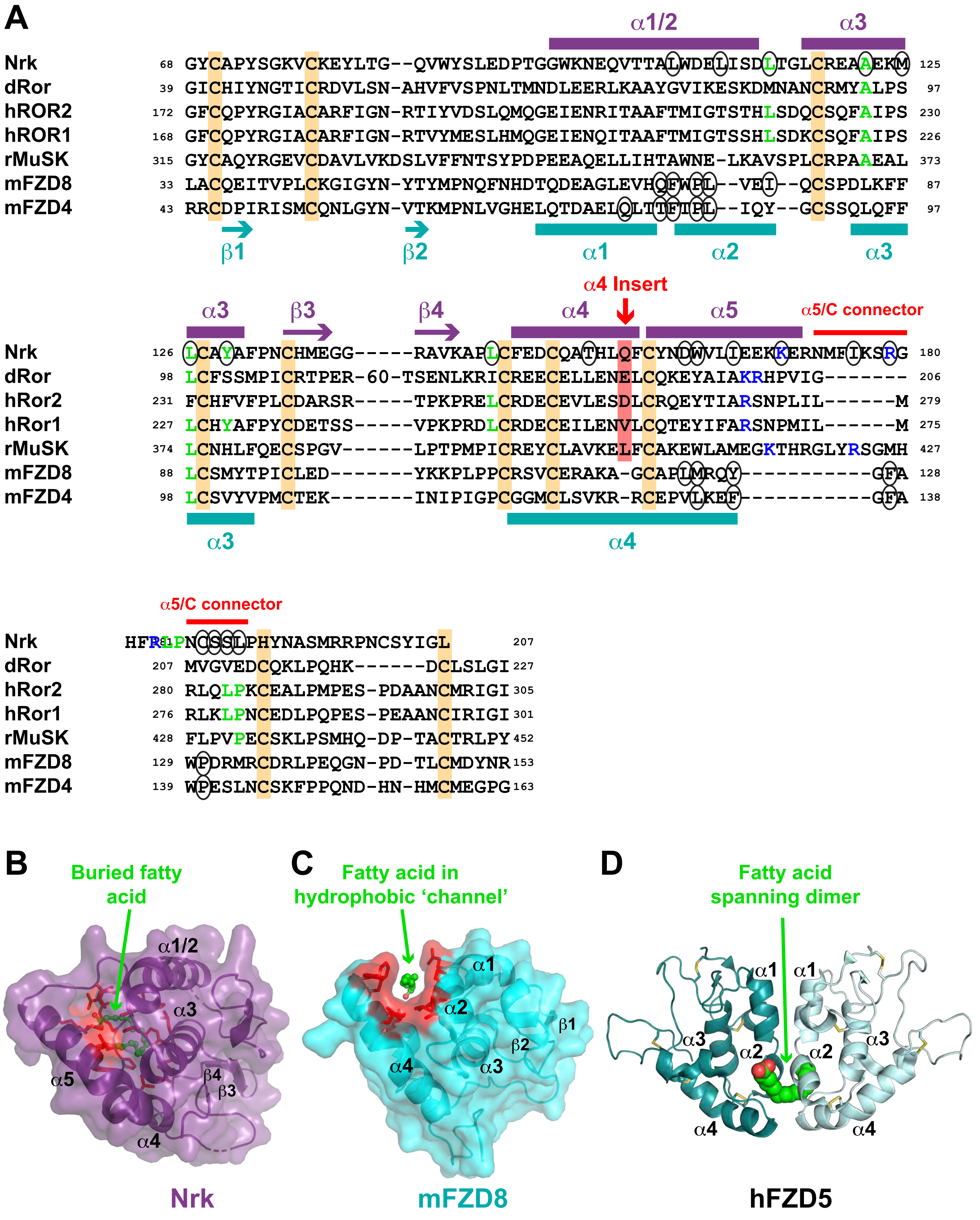

### Supplementary Figure 3

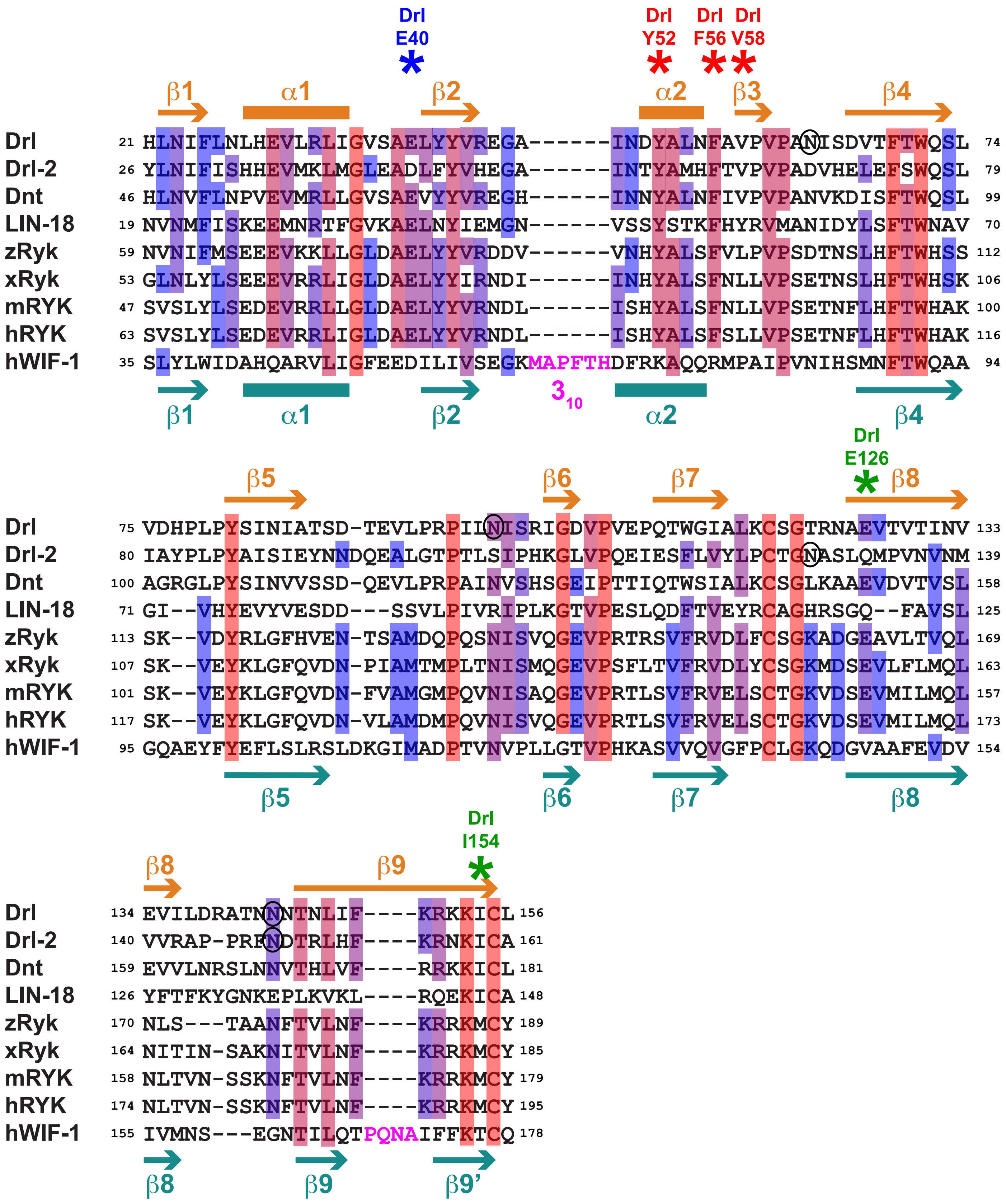

### Supplementary Figure 4

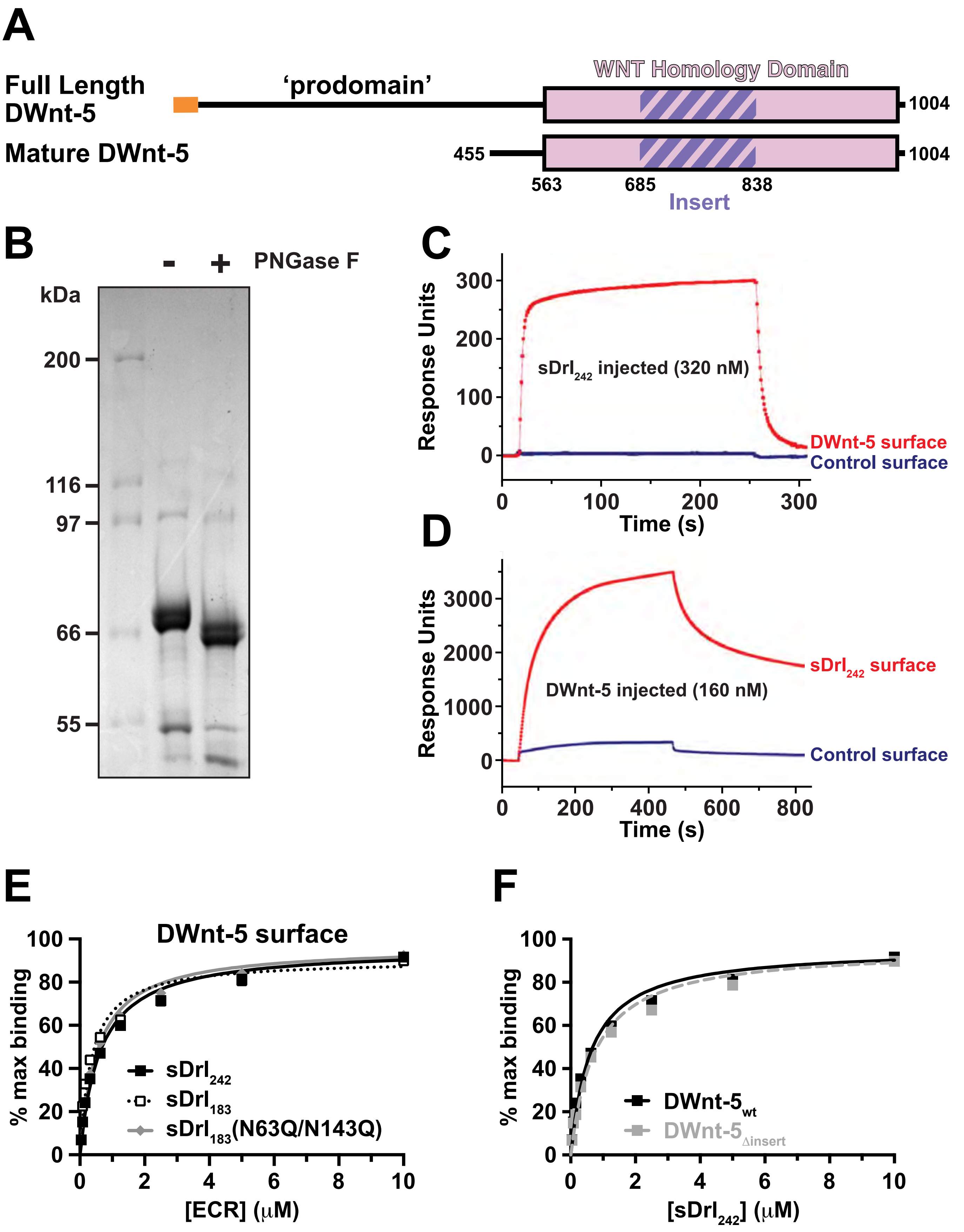

### Supplementary Figure 5

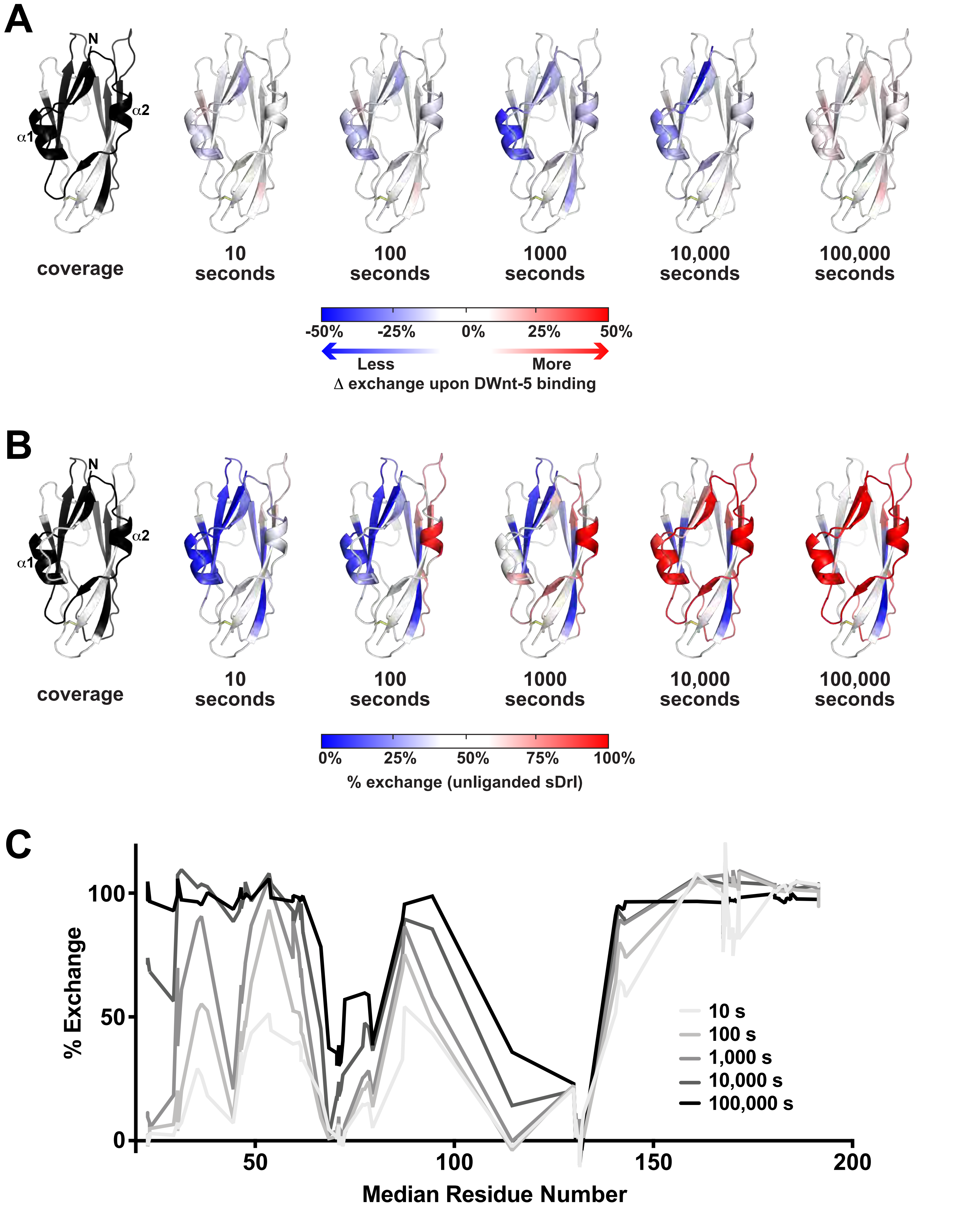

### Supplementary Figure 6

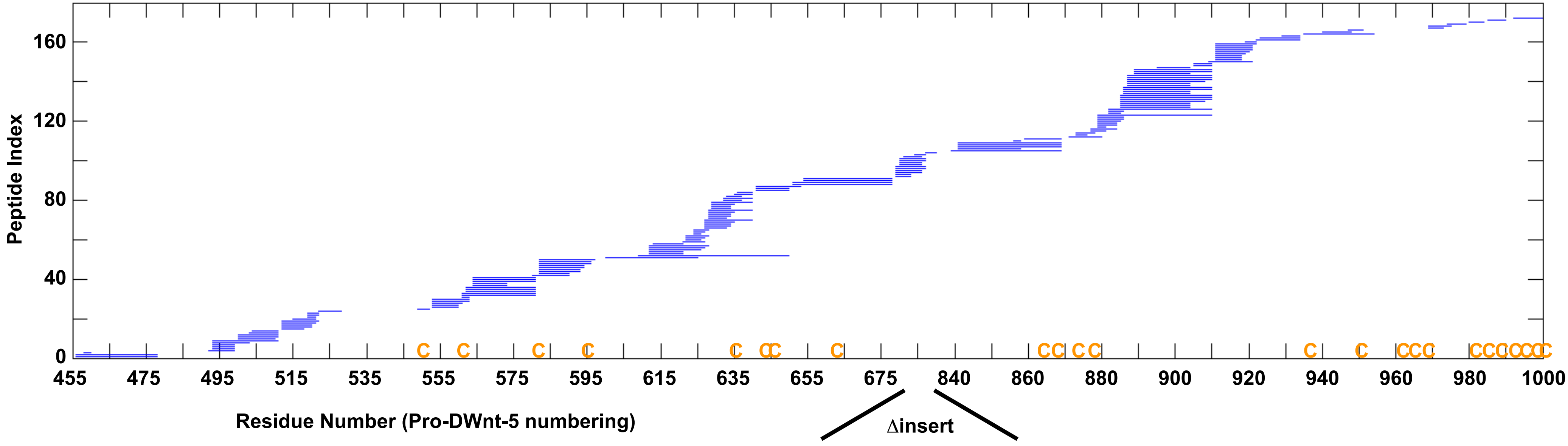
